# Structured event memory: a neuro-symbolic model of event cognition

**DOI:** 10.1101/541607

**Authors:** Nicholas T. Franklin, Kenneth A. Norman, Charan Ranganath, Jeffrey M. Zacks, Samuel J. Gershman

## Abstract

Humans spontaneously organize a continuous experience into discrete events and use the learned structure of these events to generalize and organize memory. We introduce the *Structured Event Memory* (SEM) model of event cognition, which accounts for human abilities in event segmentation, memory, and generalization. SEM is derived from a probabilistic generative model of event dynamics defined over structured symbolic scenes. By embedding symbolic scene representations in a vector space and parametrizing the scene dynamics in this continuous space, SEM combines the advantages of structured and neural network approaches to high-level cognition. Using probabilistic reasoning over this generative model, SEM can infer event boundaries, learn event schemata, and use event knowledge to reconstruct past experience. We show that SEM can scale up to high-dimensional input spaces, producing human-like event segmentation for naturalistic video data, and accounts for a wide array of memory phenomena.

## Introduction

Although sensory input arrives continuously, our subjective experience is punctuated by the perception of events with identifiable beginnings and ends (Radvansky & Zacks, 2014). These event representations are abstract and schematic, transcend specific sensory details, and allow us to generalize our knowledge across time and space. By distilling the latent structure underlying the sensory stream, event representations can be used to reconstruct the past, comprehend the present, and predict the future.

Despite the centrality of events in human cognition, there has been a notable dearth of formal models. This speaks in part to the scope and complexity of the modeling challenge; in some ways, a theory of event cognition is a theory of cognition writ large. In this paper, we take on the challenge, developing an integrated model that explains how humans segment, remember and generalize events. We frame event cognition in the language of statistical theory and argue that events serve to organize and structure continuous experience. This structure, and the generalization it affords, explains many of the empirical phenomena observed in event cognition. Along the way, we touch upon broader issues in cognitive science, including the need for structured representation, probabilistic reasoning, and parallel distributed processing. Our approach synthesizes ideas from symbolic and neural network modeling traditions, demonstrating how these ideas can work together to produce human-like competence in event cognition.

In what follows, we first summarize key empirical findings in the human event cognition literature and discuss existing theoretical treatments. We then lay out a general computational-level analysis, framing event cognition in terms of a common probabilistic generative model of events that can be inverted for different computations. This analysis forms the basic architecture of our model, which we then elaborate with a number of specific assumptions about the underlying dynamics, representations, storage and retrieval processes, and inductive biases. In the Results, we put the model to work, simulating a range of empirical phenomena. In the Discussion, we consider the strengths and weaknesses of our modeling framework, how it compares to previous theoretical treatments, and directions for future work.

## Theoretical and empirical background

A comprehensive theory of human event cognition must address the following questions:

- *Segmentation*: How do people identify event boundaries from the continuous sensory stream?
- *Learning*: How do people acquire knowledge about the internal structure of events from experience?
- *Inference*: How do people use their knowledge of events to make inferences about unobserved properties of the world?
- *Prediction*: How do people use their knowledge of events to make predictions about the future?
- *Memory*: How do people use their knowledge of events to reconstruct the past?

Here we briefly review the empirical data and theoretical ideas pertaining to each of these questions.

### Segmentation

Event segmentation has been primarily studied using subjective judgments about event boundaries in movies and text. These judgments are consistent between subjects (Newtson & Engquist, 1976; Zacks, Tversky, & Iyer, 2001) and tend to track feature changes (Hard, Tversky, & Lang, 2006) or important statistical (Baldwin, Andersson, Saffran, & Meyer, 2008; Schapiro, Rogers, Cordova, Turk-Browne, & Botvinick, 2013) and causal boundaries (Baldwin et al., 2008; Radvansky, 2012). This process appears to happen automatically and is identifiable in fMRI signals even in the absence of an explicit segmentation task (Baldassano et al., 2017; Speer, Zacks, & Reynolds, 2007).

According to *Event Segmentation Theory* (EST; Radvansky & Zacks, 2011; Zacks, Speer, Swallow, Braver, & Reynolds, 2007), people maintain an active representation of the event structure (the *current model*) that they use both to interpret the current situation and to predict what will happen within each event. When these predictions are violated, a prediction error signals the occurrence of an event boundary (Zacks, Kurby, Eisenberg, & Haroutunian, 2011; Zacks et al., 2001). This idea has been implemented in a neural network model of event segmentation (Reynolds, Zacks, & Braver, 2007), which used a context layer within a recurrent neural network to represent an event model. The activity of the context layer modulates the activity of the hidden layers of the network, thus varying the predicted dynamics by events. This architecture is similar to gating mechanisms long used in recurrent network models (Hochreiter & Schmidhuber, 1997). When simulated on a 3-D motion capture data set consisting of several short movements of a single person (e.g., chopping wood) concatenated together, prediction error in the model corresponded with transitions between events, and was used to signal updates of the context layer.

Other models of events have applied recurrent neural networks to the task of predicting the next item in a sequence. Schapiro and colleagues (2013) examined whether transition uncertainty in the environment was necessary for an event boundary. One prediction of the Reynolds model is that if a transition between two scenes is highly unpredictable then this will correspond to an event boundary. Alternatively, Schapiro and colleagues proposed that human event boundaries respect graph community structure, which can be dissociated from the predictability of scene transitions. They designed a task in which stimuli were drawn from a graph with multiple communities and, crucially, transitions between and within communities were equally probable. Thus, if people learn the underlying transition dynamics, then transitions between scenes will not generate a larger prediction error crossing a community boundary than other transitions. Nonetheless, subjects were able to identify community boundaries. Using a recurrent neural network model, they found that the network represented stimuli from the same community as more similar to each other, potentially signaling an event boundary between communities through a decrease in similarity.

A recent model by Elman and McRae (2019) considered how events are learned and play out over time for certain classes of well-defined events. They used a simple recurrent network (Elman, 1990) with localist scene representations to model the sequential dependencies that constitute an event, and probed the model’s internal representation. Trained on both toy events and human-generated sequences, the model was able to generalize between related events through co-occurrence statistics and developed meaningful internal representations.

### Learning

The process by which people learn event structure is not well understood, though several related lines of research offer some suggestive clues. One such line of research is statistical learning, proposed to explain how language learners come to delineate word boundaries in continuous speech through unsupervised observation (Aslin, Saffran, & Newport, 1998; Saffran, Aslin, & Newport, 1996) and since extended to non-linguistic and non-auditory stimuli (Fiser & Aslin, 2001; Kirkham, Slemmer, & Johnson, 2002; Saffran, Johnson, Aslin, & Newport, 1999). In a typical statistical learning task, subjects are presented with a sequence of objects and tested on successively or simultaneously presented objects without feedback. The probability distribution over sequences is potentially an important cue for learning latent temporal structure, and children and adults are able to use this structure when learning (Saffran et al., 1996, 1999). Computational models of statistical learning problems, such as word segmentation (Goldwater, Griffiths, & Johnson, 2009), feature learning (Austerweil & Griffiths, 2011) and visual chunking (Orbán, Fiser, Aslin, & Lengyel, 2008) have used Bayesian inference to group features into coherent chunks.

We contrast the temporal structured learned in auditory statistical learning tasks with relational structure, which considers symbol binding and the relationship between objects (Halford, Wilson, & Phillips, 1998; Hummel & Holyoak, 2003; Kemp, Tenenbaum, Niyogi, & Griffiths, 2010). Arguably, visual statistical learning requires relational structure (Austerweil & Griffiths, 2011), but temporal and relational structure are nonetheless distinct. Events are thought to contain both temporal and relational structure (Tversky, Zacks, & Hard, 2008), and we would expect this structure to be unpredictable from low-level features (Richmond & Zacks, 2017). Bayesian models have been developed to explain relational structure learning (Kemp & Tenenbaum, 2008; Kemp et al., 2010), and we develop a variant of these models for learning event schemata.

Neural networks also provide convenient models for learning the types of sequential dependencies that constitute events. Typically, the sequential dependencies that constitute an event are modeled with recurrent networks (Elman, 1990; Hochreiter & Schmidhuber, 1997), which have strong theoretical guarantees in terms of what they can learn (Siegelmann & Sontag, 1991, but see Geman, Bienenstock, and Doursat 1992), and have been empirically effective in computational domains with long-term sequential dependencies, such as language and reinforcement learning (LeCun, Bengio, & Hinton, 2015). Because neural networks rely on distributed representations, they provide a basis for representing conceptual similarity though vector similarity (Hinton, McClelland, & Rumelhart, 1986), and are well suited for smooth generalization. These distributed representations can in some cases support symbolic structure via algebraic manipulation (Doumas & Hummel, 2005; Hummel & Holyoak, 2003; Plate, 1995; Smolensky, 1990), which allows them to encode the higher-order symbolic processes that constitute events.

### Inference

Event structure can inform inferences people make about parts of their environment they did not experience. As we move through our natural world, we do not always have immediate perceptual access to relevant features of our immediate surroundings. For example, if a person hears a door open behind them, they can make several inferences about the current scene that are relevant to the ongoing event, even though they do not have full perceptual access to each feature.

Inferences about events have historically been investigated with reading and memory tasks in which a subject reads a narrative and either reading time (Altmann & Mirković, 2009; McKoon & Ratcliff, 1992) or memory (Bower, Black, & Turner, 1979; Bransford, Barclay, & Franks, 1972) is used as a probe for their inferences about unstated aspects of the event. One of the main themes in the reading comprehension literature is that only a subset of inferences are made online, specifically those that are necessary for satisfying some notion of coherence in the text, although there is disagreement about what exactly this entails (Graesser, Singer, & Trabasso, 1994; McKoon & Ratcliff, 1992; Trabasso & Van Den Broek, 1985). In the constructionist account (Graesser et al., 1994), elements of the event model are used to fill in aspects of the event when they are likely to be relevant, and these inferred portions of the event are more quickly accessed and less surprising when they subsequently occur. This implies an averaging effect where inferences about the unstated aspects of an event reflect commonly experienced configurations. This is conceptually similar to adaptive statistical biases seen in other domains, such as the estimation of spatial location (Huttenlocher, Hedges, & Duncan, 1991).

### Prediction

Event structure shapes predictions about the future. This can be seen in serial reaction time tasks, in which subjects respond to cues more quickly when they are generated from repeated, predictable patterns than when they are generated as a random sequence (Nissen & Bullemer, 1987). This form of prediction has been argued to be implicit (Reber, 1989; Robertson, 2007) and is consistent with earlier ideas that people learn dynamic schemata such as “scripts” to guide their actions (Lashley, 1951; Schank & Abelson, 1977).

There is also a long history of studying prediction in language comprehension. While reading, subjects fixate less often and for shorter durations on highly predictable words (Ehrlich & Rayner, 1981) and are slower to process unexpected words (Schwanenflugel & Shoben, 1985). Words completing sentences in a nonsensical way elicit an N400 event-related potential, thought to signify a surprising or unexpected stimulus (Kutas & Hillyard, 1984). Event knowledge specifically appears to influence language comprehension (McRae & Matsuki, 2009); individual words cue event-based knowledge (Altmann & Kamide, 1999; Hare, Elman, Tabaczynski, & McRae, 2009; McRae, Hare, Elman, & Ferretti, 2005), and combinations of words narrow the scope of perceived events (Matsuki et al., 2011).

Neural measures also offer support for the proposal that people continually leverage sequential structure to make online predictions about upcoming experiences (Cohn, Jackendoff, Holcomb, & Kuperberg, 2014; Schiffer & Schubotz, 2011). These predictions are influenced by event structure. Behaviorally, people make better predictions within an event than across event boundaries (Zacks et al., 2011), and unpredictability across event boundaries is associated with prefrontal cortex, striatum, and hippocampus (Axmacher et al., 2010; Lisman & Grace, 2005; Ranganath & Rainer, 2003; Zacks et al., 2011). These findings are consistent with the hypothesis that failure in prediction is used to signal event boundaries (Reynolds et al., 2007; Zacks et al., 2007).

### Memory

Memory can be both aided and impaired by knowledge of event structure. Event boundaries induce selective trade-offs in memory that depend on the exact study design and memory measure. Sequential recall and temporal order memory are worse across event boundaries than within events (DuBrow & Davachi, 2013, 2016; Heusser, Ezzyat, Shiff, & Davachi, 2018), but memory for specific items has been found to be higher at event boundaries (Heusser et al., 2018), and the presence of an event boundary can increase overall recall (Pettijohn, Thompson, Tamplin, Krawietz, & Radvansky, 2016). Moreover, individuals with better event segmentation ability perform better on subsequent memory tests (Sargent et al., 2013; Zacks, Speer, Vettel, & Jacoby, 2006). An ongoing event appears to have a privileged role in memory as well, as memory for items within an ongoing event is often better than immediately following an event boundary in a way that is not recovered by returning to the original context (Radvansky & Copeland, 2006; Radvansky, Krawietz, & Tamplin, 2011).

The *Event Horizon Model* (Radvansky, 2012; Radvansky & Zacks, 2014) offers a conceptual explanation for these findings. According to this model, people track the causal structure of events and use this structure to aid memory retrieval. This causal structure leads to better memory for items overall but can lead to memory interference under certain conditions, such as when recall depends on retrieval of a single event model and the presence of multiple events can introduce noise. The Event Horizon Model further hypothesizes that working memory maintains privileged access to the current model, which thus results in better memory retrieval. According to this account, sequential recall might be impaired across event boundaries because it relies on two competitive event models, whereas overall recall would be improved because it is non-competitive.

Event knowledge has also been implicated in false memory paradigms (Bower et al., 1979; Bransford et al., 1972). In these paradigms, subjects report remembering unstated details of a story that were nonetheless consistent with the narrative. This effect is parametric, such that more experiences with similar stories increase script-consistent false memories (Bower et al., 1979). This suggests that people use event knowledge to fill in the gaps of their memory and are consistent with *script theory* in which people organize sequential processes in terms of an abstract schema, or script, that organizes perception and influences memory (Schank & Abelson, 1977). Conceptually, this is similar to the idea of *pattern completion*, where a partial memory trace is reconstructed with reference to learned pattern of activity (McClelland, McNaughton, & O’Reilly, 1995; Norman & O’Reilly, 2003), and may be adaptive if it facilitates inference about unobserved aspects of an event.

## Limitations of previous theoretical accounts

Most previous theoretical accounts of event cognition have been non-computational (Radvansky, 2012; Radvansky & Zacks, 2014; Zacks et al., 2007). While these accounts provide valuable theoretical insight into event perception and memory, they do not offer the same level of detailed prediction as a computational model. Computational accounts of events have typically focused on event segmentation and learning the sequential dependencies within events, often using recurrent neural networks. A model by Reynolds et al. (2007) proposed that recurrent neural networks could be used to learn event dynamics and event boundaries using prediction error as a signal for both. In this account, event boundaries are driven by prediction error, or a discontinuity in what was expected by the event model and what was observed. This model was challenged by Schapiro et al. (2013), who argued that the Reynolds model implied event boundaries driven by environmental unpredictability. Schapiro and colleagues demonstrated empirically that subjects delineate event boundaries reflecting underlying community structure even when equated for uncertainty. They further showed that a recurrent neural network would implicitly learn this community structure in the absence of an explicit segmentation mechanism and argued against the prediction error account.

While the Reynolds et al. (2007) and Schapiro et al. (2013) models provide competing accounts of event segmentation, neither model fully explores the generalization of event dynamics nor incorporates structure into their representations. This limits their ability to explain data on script memory and text comprehension, and it has been further argued that structured event representations are a better account of human generalization (Goodman, Ullman, & Tenenbaum, 2011; Kemp & Tenenbaum, 2008; Richmond & Zacks, 2017).

A recent model by Elman and McRae (2019) proposes a simple recurrent network as a model of event dynamics and examines how the representation in the network changes across time in a toy structured world. Interestingly, the model can infer missing fillers in their appropriate roles based on historical co-occurrence statistics. When the model is trained on multiple event types, it can generalize between them by leveraging a shared representational space. This model uses localist representations of symbolic structure, which may not scale to naturalistic datasets. Furthermore, as the model is concerned with learning the internal structure of events, there is no explicit mechanism to identify an individual event or determine its boundaries.

In all of these computational models, events are modeled as dynamical processes that unfold over time. To our knowledge, how these processes interact with memory has not been addressed with computational modeling. The *Temporal Context Model* (Howard & Kahana, 2002) and the related *Context Maintenance and Retrieval* model (Polyn, Norman, & Kahana, 2009) treat memory as a dynamical process in which an evolving context induces a temporal organization in memory. These models suggest dynamics similar to what we would expect when learning events but do not provide a mechanism for partitioning events. Related models of memory support some forms of inference (e.g., Nelson & Shiffrin, 2013; Shiffrin & Steyvers, 1997), but these models tend not to incorporate the dynamical processes that define events.

Finally, no prior computational model of event cognition has attempted to explain naturalistic data (e.g., real videos). While the use of naturalistic data is not a theoretical challenge to any prior model of event segmentation, it is nonetheless an important practical consideration. To validate our computational theories of cognition, we should strive to show that they scale to real-world problems. Otherwise, it is not clear whether our models of cognition work only when constrained to artificial tasks. More broadly, while several computational models have addressed parts of event cognition; to our knowledge, no model has attempted to address all aspects of the problem.

## The Structured Event Memory model

We propose a model, *Structured Event Memory* (SEM), that systematically addresses the 5 desiderata for understanding event cognition (segmentation, learning, inference, prediction, memory), overcoming the limitations of prior models.

Our model is a computational-level analysis of event segmentation and memory that frames these tasks in terms of probabilistic reasoning. At a high level, our model assumes that people organize *scenes* into *events* that are defined by a shared temporal structure. People use the temporal structure of events to organize their perception, simultaneously learning this structure while using it to predict. We further propose that people use this learned event structure to aid reconstructive memory by compensating for information loss with the predictable temporal structure of events.

The model has several key computational mechanisms. First, we use a distributed representation to encode the structure of individual scenes. By scenes, we mean a description of the environment containing a relevant set of objects and the relations between them (similar to a “situation” in a situation model). For example, for a person watching a movie, a scene might be the experience of sitting in a dark room holding popcorn, or it might describe the arrangement of characters on the screen and its setting. In the context of a psychological study, a scene might describe stimuli currently on a screen and the context of sitting in front of a computer.

We model events as stochastic dynamical processes over these scenes. Each event token is a sample from this process, associated with a sequence of scenes. The learned event dynamics are used to organize scenes into distinct event types (analogous to a *script* or *event schema*) via a clustering process where the statistics of similar event tokens are pooled for the purpose of learning and generalization. Finally, we assume these learned structures are used to improve a capacity limited, and thus noisy, memory by regularizing noisy memory traces towards prototypes defined by each event type.

Each of these computational mechanisms has a different mathematical instantiation, and their interlocking behavior is key to our normative account of events (Figure 1). To embed the structure of individual scenes as vectors, we use holographic reduced representations (Plate, 1995), allowing us to encode logical structure while maintaining the advantages of distributed systems. We model event dynamics with a recurrent neural network defined over the holographic embeddings. Scenes are assigned to particular events via a Bayesian clustering process. Event dynamics and clustering scenes into events are two distinct forms of temporal structure that complement each other. Clustering scenes into events allows for rapid generalization and efficient credit assignment whereas learning the dynamics of events with recurrent neural networks represents an internal temporal structure within each event. Memory is instantiated through a form of Bayesian smoothing, in which a set of noisy memory traces is combined with learned event dynamics in a probabilistic reconstruction process. We now discuss these components in more detail.

**Figure 1.**
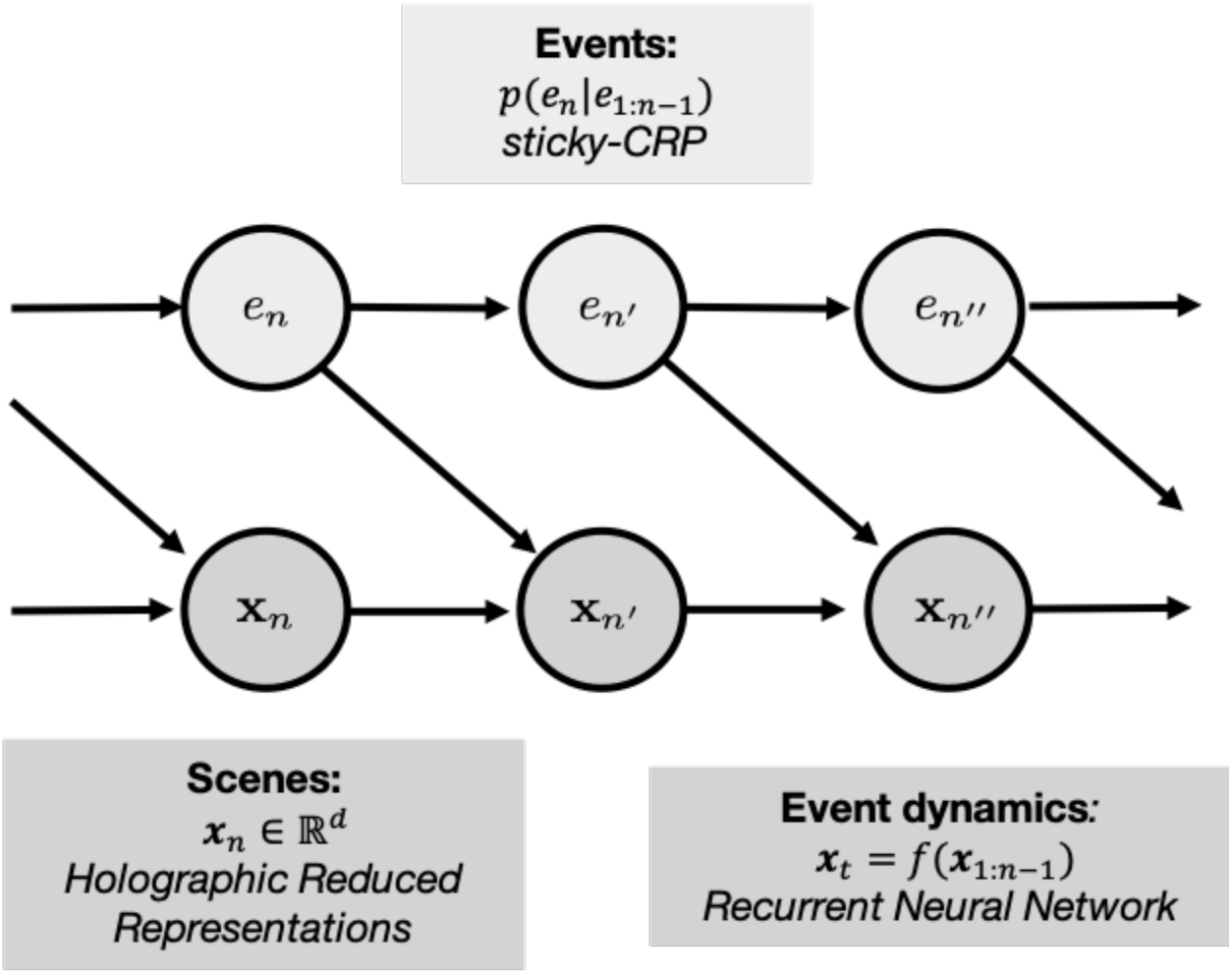
Generative model of events. Events are assumed to follow their own evolution, defined by a sticky Chinese Restaurant Process (sticky-CRP), and to constrain the dynamics of scenes. These scenes combine both structured and unstructured information in distributed representations using holographic reduced representations (HRRs). Recurrent neural networks are used to learn the non-linear dynamical system that defines the dynamics within individual events.

**Figure 2.**
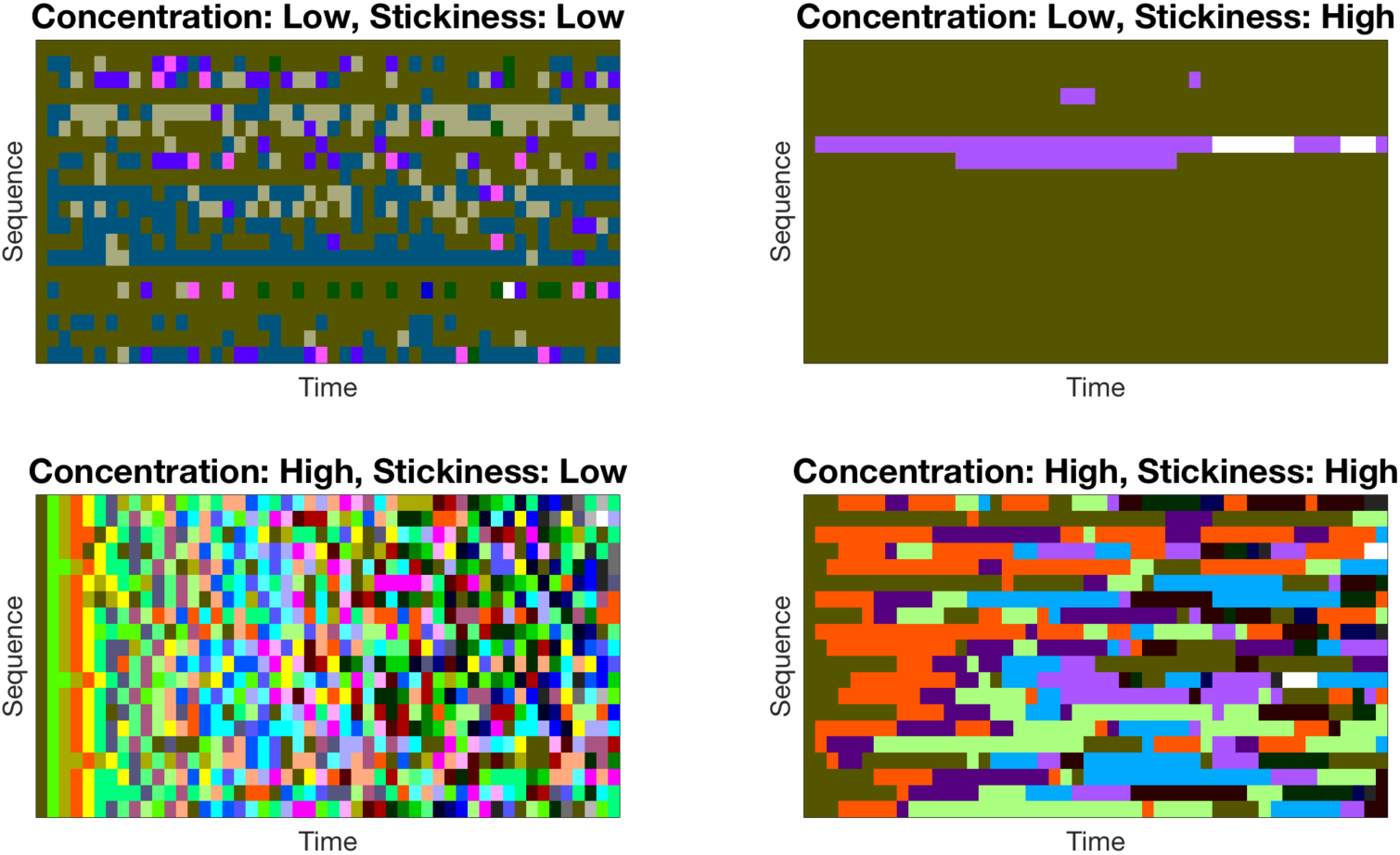
Samples from the generative process of events under different parameter regimes. Each row in each panel is a single draw from the process across time. Colors indicate when different event schemata are active. High values of the concentration parameter *α* (bottom row) lead to more unique events. High values of the stickiness parameter λ (right column) lead to longer event durations.

### A generative model of events

We make the following assumptions about events:

1. Humans do not know *a priori* how many distinct event types there are; this must be discovered from the data.
2. Humans attempt to reuse previously learned event models whenever possible. This constitutes a simplicity bias in the sense of Ockham’s razor, which facilitates generalization between events.
3. Events are temporally persistent, such that there is a high probability that any two consecutive moments in time are in the same event.
4. Events define dynamical systems, generating predictions of successor scenes as a function of the current scene.
5. Events have latent structure: each instance of an event (and the scenes within events) may be unique with respect to specific percepts but nonetheless shares a latent structure (such as the relationships between objects) with similar events.
6. Events are used in memory retrieval to regularize noisy memory traces. Event knowledge can compensate for missing or corrupted information in a memory trace by “filling in the gaps” using knowledge about event structure.

Assumptions 1, 2 and 3 stipulate constraints on the generative process that produces observed scenes. We now describe how these constraints can be translated into a mathematical model.

A simple Bayesian nonparametric process known as the sticky Chinese restaurant process satisfies the first three assumptions (sticky-CRP; Fox, Sudderth, Jordan, & Willsky, 2011; Gershman, Radulescu, Norman, & Niv, 2014).^1^ The sticky-CRP is a generative process that sequentially assigns time points to events according to past event frequency and recency while, as a nonparametric process, maintaining some probability that a new event will be generated at each moment in time (cf. assumption 1). The sticky-CRP and related processes are commonly used as the prior in non-parametric clustering algorithms, in which they function as a probability distribution over partitions (Fox et al., 2011). Later, we will invert our generative model of events for the same purpose and use the sticky-CRP as a prior over the assignment of scenes into events.

Under the sticky-CRP, higher-frequency events are more likely to be repeated than low-frequency events, and events are likely to be repeated sequentially. Formally, at time *n* the next event *e_n_* is drawn from the following distribution:

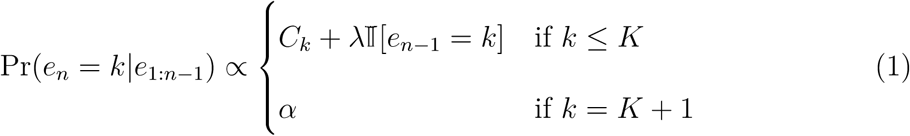

where *e_n_* is the event assignment, *K* is the number of distinct event types in *e*_1:_*_n_*_−1_, I[·] = 1 if its argument is true (0 otherwise), and *C_k_* is the number of previous timepoints assigned to event k. The concentration parameter *α* > 0 determines the simplicity bias (cf. assumption 2); smaller values of *α* favor fewer distinct events. The stickiness parameter λ ≥ 0 determines the degree of temporal autocorrelation (cf. assumption 3); higher values of λ favor stronger autocorrelation.

The remaining assumptions relate to the nature of event schemata (assumptions 4 and 5) and memory retrieval (assumption 6). In the following sections, we discuss how these assumptions can be instantiated in a computational model. First, we discuss the representational space in which event models, as dynamical systems, operate (cf. assumption 4). Representation is critical for generalization, and to that end, we discuss a structured representation in vector space that facilitates generalization via continuous functions. We then discuss event dynamics and how they are learned, how event segmentation occurs in the model. Finally, we present our model of event memory.

### Scene representations

We assume that event schemata define dynamical processes over scenes, in which the event model is used to generate a prediction about the next scene given the recent history of scenes. Formally, we define a function that takes in a scene *s* ∈ *S* and returns a successor scene *s*’. We further assume that for each scene *s*, there exists a distributed (vector) representation **x** ∈ ℝ*^d^*. Whereas a scene can be thought of as a ground-truth description of the external world, **x** can be thought of as a representation of the features of *s* relevant to an agent.

A core assumption is that the vector representation of the scene is distributed and encodes features in a similarity space (Goodfellow, Bengio, & Courville, 2016). For example, if *cat* and *dog* have meaningful representations, they may share several features encoded in the space, such as *isSmall* and *isFurry*. As pure symbols, *cat* and *dog* are as distinct from each other as any other pair of symbols. As a consequence of this similarity space, similar scenes have similar vectors such that the vector representation of *dog* is close in Euclidean space to the vector representation of *cat*.

A primary motivation for using distributed representations is that they facilitate smooth generalization. As we will discuss in the next section, defining scenes in vector space allows us to parametrize event dynamics over arbitrary scenes. We will assume that these event dynamics are smooth, such that if *f* is a function that represents the event dynamics and **x** ≈ **y**, then *f* will generally have the property *f* (**x**) ≈ *f* (**y**) (Goodfellow et al., 2016). An embedding space that encodes similar scenes with similar vectors will facilitate this type of generalization naturally with parameterized functions; by contrast, were we to use a tabular representation, we would have to define transitions over an intractably large discrete space that does not permit smooth generalization.

Distributed representations have the further advantage that they are representationally compact. A purely symbolic representation encodes a dimension for each binary feature, whereas a distributed representation can take advantage of the correlational structure to encode scenes in a lower-dimensional space (Goodfellow et al., 2016). This is related to dimensionality reduction, in which a representation is projected into a low-dimensional space that preserves relevant features and discards irrelevant features. This property is important for computational reasons, as it is computationally intractable to estimate dynamical systems in high dimensional spaces, and event dynamics may be more easily learnable in an appropriately chosen low-dimensional representation (Richmond & Zacks, 2017). As a practical matter, we need a principled way to construct a low-dimensional scene representation to model naturalistic data sets. Later in the paper we will present simulations with video data and show that convolutional neural networks are well suited to this task for visual domains (Fukushima, 1980; Kietzmann, McClure, & Kriegeskorte, 2018; LeCun et al., 2015). Low-dimensional representations can be learned in visual data with unsupervised methods (Hou, Shen, Sun, & Qiu, 2017; Radford, Metz, & Chintala, 2015). In our simulations, we use a variational autoencoder (VAE; Kingma & Welling, 2013), an unsupervised convolutional neural network, to learn the representational space (see Appendix A for details). We note that while the representation of scenes is important for our theoretical model, we are agnostic to the specific details of how it is learned in the brain.

In addition to these unstructured features of scenes, we further assume that scenes encode relational structure. By this, we mean that scenes contain relational information about how particular fillers are bound to particular roles (Radvansky & Zacks, 2011). For example, if a person is holding a phone, the scene would contain not just a reference to the objects *person* and *phone*, but also a specific binding between the objects and their roles in the relationship *holding*. A faithful vector representation of structured scenes would contain this structure when relevant. There are several ways binding can be implemented in vector spaces, such as using tensor products (Smolensky, 1990), holographic reduced representations (Plate, 1995), or binding by synchrony (Hummel & Holyoak, 2003). To be clear, we are not claiming that symbolic computation is necessary for all event representations. Undoubtedly, unstructured (non-relational) features play an important role. One might expect, for example, that a sudden change in the amount of ambient light might correspond to an event boundary. The importance of these unstructured features partially motivates encoding symbolic structure in distributed representations, as it allows us to combine both in a single scene representation.

In this paper, we will focus on holographic reduced representations (HRRs), a member of a family of representations known as vector symbolic architectures (Gayler, 2004). Unlike scalar multiplication, the tensor (outer) product of two vectors is not commutative, such that the tensor product of two vectors encodes order information. This property is sufficient for a binding operation, but tensor products do not scale well with multiple terms. Vector symbolic architectures get around this problem by using a compressed form of the tensor product as a binding operation. The HRR, for example, uses circular convolution. In addition, independent terms can be combined with vector addition, thus allowing a scene with multiple bound terms to be represented with a vector of fixed dimensionality, regardless of the complexity of the underlying formula.

Concretely, consider the expression “dog chases cat.” This takes the logical form *chase(dog, cat)*, where dog occupies the agent role and cat occupies the patient role. If we have vectors representing both roles and fillers (where our fillers are *dog*, *cat* and *chase* and our roles are *agent*, *patient* and *verb*), then the scene vector is constructed according to:

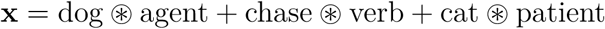

where ⊛ denotes circular convolution. The underlying components can be approximately decoded using circular correlation, a simple algebraic operation, directly from the composed scene vector (see Plate, 1995, for details).

The HRR is convenient because both the encoding operation and approximate decoding operation can be accomplished by efficient algebraic manipulations. Furthermore, the operations of HRRs preserve the similarity of the composed terms (see Appendix B for details), meaning that structural properties can be encoded as features of the embedding space. For example, we can decompose a term as a linear combination of structured and unstructured features, such as

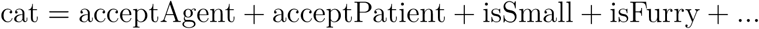

where the features *acceptAgent*, *acceptPatient* correspond to valid relational roles *isSmall*, *isFurry* are unstructured features. If the term *dog* corresponds to a similar vector with shared composed features, then a scene composed with *dog* will be similar to a scene composed with *cat*. Concretely, the scene *chase(cat, mouse)* will be close in vector space to the scene *chase(dog, mouse)* because of the similarity of the fillers. We assume that each unique object is represented as the same vector across multiple scenes. For example, two different cats would be represented with two different (but similar vectors) whereas a single cat would be represented across time with a single vector. This encoding scheme also allows properties of an individual token to change, for example, if a cat is dry in one scene and wet in the next, the vector representation of the dry cat can be modified to encode this new property by adding a new feature “isWet” through vector addition. These representations would be similar but nonetheless distinct, representing the change in the token over time. As noted above, this representation facilitates smooth generalization of event dynamics by parameterized functions. Finally, it is worth noting that the composed terms of an HRR are not precisely equivalent to a logical representation, but are approximations, as the circular convolution necessarily loses information to preserve dimensionality. In practice, these concerns are minor, as the capacity of HRRs to encode unique symbols is quite high (Eliasmith, 2013) and can be augmented through permutation of the underlying terms (Kelly, Blostein, & Mewhort, 2013).

An important question that we have not addressed is how structured representations can be learned. For the purpose of the current work, we are largely concerned with how structured vector spaces are important in event cognition and are not making strong commitments as to how they are learned. We assume these representations where noted and construct them by drawing Gaussian random vectors for each independent feature, conjoining these features with vector addition. In the *cat* example above, we would draw the random vectors for all of the features *acceptAgent*, *acceptPatient*, *isSmall*, and *isFurry* and compose them with vector addition to create the full symbol *cat*. This creates the desired vector space in which symbols have independent linear components. Similar predicate representations are common in connectionist models of semantic cognition (Rogers & McClelland, 2005). Moreover, vector spaces with independent linear components are learnable with fully unsupervised methods, as both word (Mikolov, Sutskever, Chen, Corrado, & Dean, 2013) and visual scene embeddings (Radford et al., 2015) often have this property.

### Event dynamics

We model event schemata as dynamical systems over scenes: the probability of observing a successor scene *s_n_*_+1_ given the current scene *s_n_* is conditioned on the event *e_n_* such that Pr(*s_n_*_+1_|*s_n_, e*), where *s_n_* is the scene at time index *n*. In SEM, we assume this probability distribution is defined with a smooth function *f* over the embedded scenes and parameterized by *θ_e_*, such that

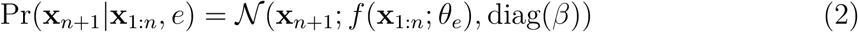

where *β* is a vector of noise variance parameters (one for each feature dimension), and **x***_n_*_+1_ and **x**_1:_*_n_* are the vector embeddings for the successor scene *s_n_*_+1_ and all of the previous scenes, respectively. As a consequence of this formulation, scenes that are highly similar to the prediction of an event schema have high probability under the schema, naturally allowing for smooth generalization.

We are agnostic to the shape of the function *f* and the form of the parameters *θ_e_*. We only require that *θ_e_* is learnable from observations, and we assume that *θ_e_* is a set of parameters unique to each event. In our simulations, we model *f* as a recurrent neural network (Figure 3), and *θ_e_* is a set of weights and biases that determine the activity of the network. At each time-step, the network takes the vector embedding for the current scene, **x***_n_*, as an input, and generates a prediction of the successor scene, **x***_n_*_+1_ as its output. The network also maintains an internal state that is updated on each time-step and that used when making its prediction.

**Figure 3.**
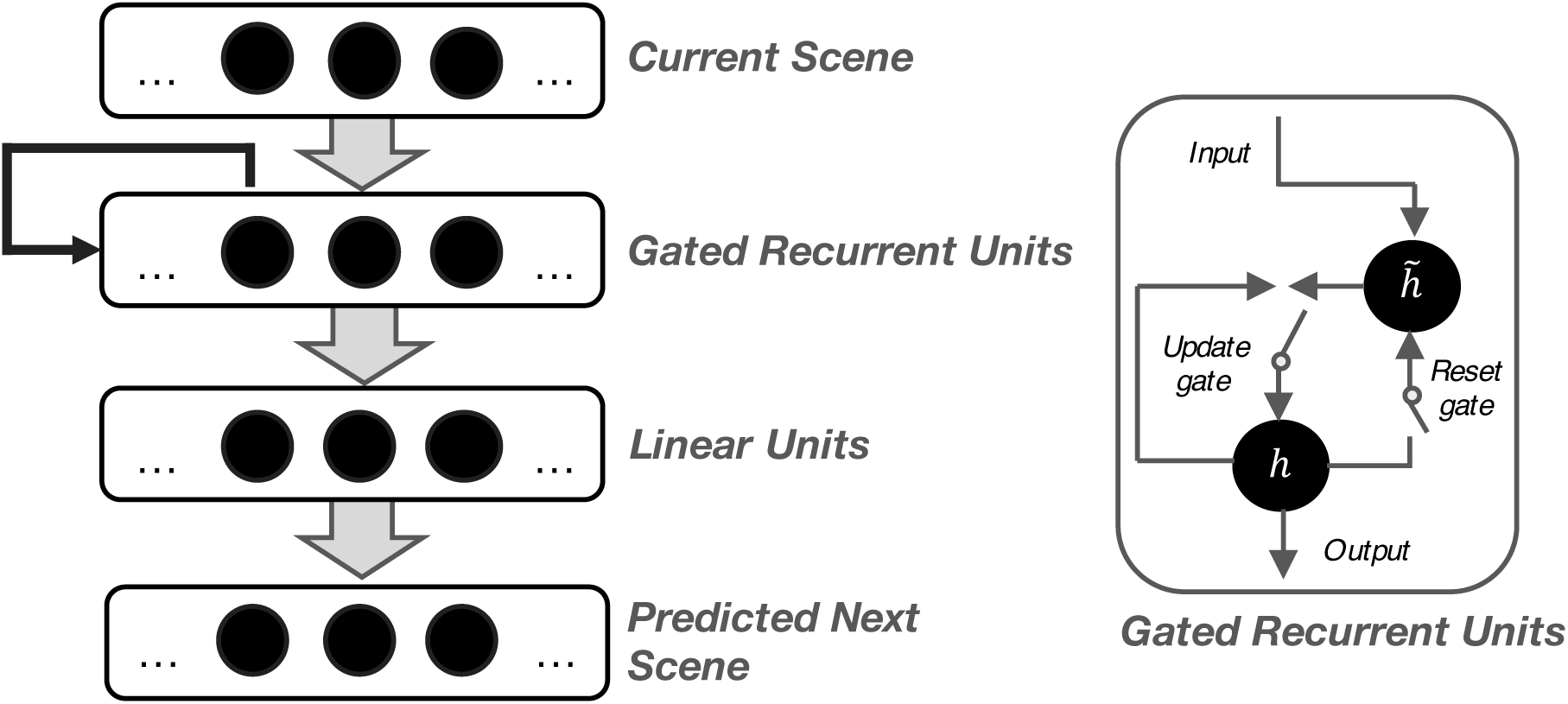
Event Dynamic Neural Network. A four-layer network took in the current scene as the input and predicted the next scene as the output. The network has two hidden layers: one with gated recurrent units (GRU; Cho et al., 2014) and a leaky rectified linear output and a second layer of fully connected, linear units. GRUs provide recursion by maintaining an internal state of the network. This state is controlled by two gates, an *update* gate and a *reset* gate, which control the degree to which the hidden layer is updated or reset with new input, respectively.

Except where noted, we used a fully connected, four-layer network with gated recurrent units (GRUs; Cho et al., 2014) with a leaky rectified linear activation function (*α* = 0.3; Maas, Hannun, & Ng, 2013) as a non-linearity and 50% dropout for regularization (Srivastava, Hinton, Krizhevsky, Sutskever, & Salakhutdinov, 2014). The GRU has two gates that control its recurrent activation, an *update* gate that controls whether the current hidden state is updated with new information and a *reset* gate that controls whether to use the previous hidden state in this update. We have chosen the GRU because it is a simplified variant of the Long-Short Term Memory network (Hochreiter & Schmidhuber, 1997) that is computationally less demanding to train but nevertheless performs well on natural datasets (Chung, Gulcehre, Cho, & Bengio, 2014). Gating networks maintain an internal state that the network learns to update (or gate) through backpropogation (Figure 3, right). This internal state allows the network to maintain a representation of previous stimuli, which it can use to earn long-term sequential dependencies (Hochreiter & Schmidhuber, 1997) and problems that require memory (O’Reilly & Frank, 2006). Reynolds et al. (2007) previously found that gating networks were better able to learn event structure than feed-forward networks. A second, non-recurrent hidden layer with a linear output is used to produce the output. The networks were implemented in Keras (Chollet et al., 2015) and were trained with batch updates of cached observations using the Adam stochastic optimization algorithm (Kingma & Ba, 2014). Parameter values for this optimization are listed in Table 2.

**Table 1.** Model Parameter values. The parameter values for each simulation are listed separately in each row.

| Simulation | Embedding Dim. | $\nu_0$ | $s_0^2$ | $\lambda$ | $\alpha$ | $\beta$ | $\tau$ | $\epsilon_e$ |
| --- | --- | --- | --- | --- | --- | --- | --- | --- |
| Zacks et al. (2006) | 100 | 10 | 0.06 | $10^4$ | 0.1 | - | - | - |
| Schapiro et al. (2013) | 25 | 1 | 1.0 | $10^5$ | 0.01 | - | - | - |
| Structured Event Boundaries | 266 | 100 | 0.305 | 1.0 | 1.0 | - | - | - |
| Bower et al. (1979) | 185 | 10 | 0.24 | 1.0 | 10.0 | 2 | 0.15 | 0.75 |
| Radvansky and Copeland (2006) | 25 | 1 | 0.2 | 10.0 | 1.0 | 2 | 0.1 | 0.25/0.75 |
| Pettijohn and Radvansky (2016) | 25 | 1 | 0.2 | 10.0 | 1.0 | 2 | 0.1 | 0.25 |
| DuBrow and Davachi (2013, 2016) | 25 | 1 | 0.2 | 10.0 | 1.0 | 2 | 0.1 | 0.25 |

**Table 2.** Neural network optimization parameter values. Parameter values for the Adam optimization algorithm used to train the recurrent neural network used to estimate event dynamics. Each simulation used the same optimization parameters.

| | learning rate | $\beta_1$ | $\beta_2$ | $\epsilon$ | decay | n_epochs |
| --- | --- | --- | --- | --- | --- | --- |
| Parameter | 0.01 | 0.9 | 0.999 | 1e-08 | 0.0 | 10 |

We chose a recurrent network so that the event schema would be sufficiently flexible to learn the event dynamics. Previous theoretical modeling suggests that event dynamics can be represented with recurrent neural networks (Elman & McRae, 2019; Reynolds et al., 2007; Schapiro et al., 2013), and similar recurrent neural networks are thought to be biologically plausible (O’Reilly & Frank, 2006). Furthermore, Reynolds et al. (2007) found that recurrence improved learning event dynamics when compared to otherwise identical feed-forward networks. Nonetheless, we do not believe that any of these specific implementation details are critical, and other variants may make similar predictions.

The noise over scene transitions in equation 2 is assumed to be Gaussian with a diagonal covariance matrix, parametrized by the vector of noise parameters 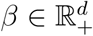. This diagonal covariance matrix allows us to model the noise of each dimension independently, while still allowing this noise distribution to be learnable. In contrast, a spherical covariance matrix could result in a single dimension driving segmentation and estimates of the full covariance matrix of a high-dimensional Gaussian are not reliable in small sample sizes due to the curse of dimensionality. The vector *β* is estimated using maximum a posteriori estimation, assuming an inverse-χ^2^ prior parametrized by ν degrees of freedom and a scale of *s*^2^ (Gelman et al., 2013).

We separately define an initial condition *f*_0_ for the function *f*, which is estimated from the data with a uniform prior over *f*_0_. The transition probability for scene *s_t_*_+1_ following an event boundary is thus:

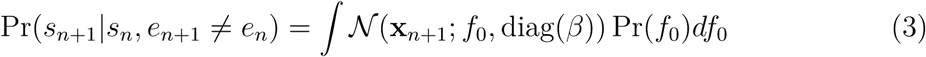

Importantly, this initial condition probability is different for experienced and novel events. The prior over *f*_0_ is important for novel events, as it allows us to define a scene transition probability for an unseen event by integrating over the prior. When the event *e_n_*_+1_ is a previously unseen event, *e_new_*, the transition probability in equation 3 reduces to a constant. For experienced events, we simplify this probability function by ignoring the prior Pr(*f*_0_) and instead use a point estimate of *f*_0_. As we discuss in the following section, we use the probability Pr(*s_n_*_+1_|*s_n_, e_new_*) to infer the event boundaries and therefore require a definition of this term for unseen events.

### Event segmentation

Having defined the generative process for events, the representational space for scenes, and the scene dynamics, we can now pose questions for the computational model. A primary challenge for the model presented above is how it can learn and segment events without an external training signal. We define event segmentation as the process of assigning an event label to each scene, and in terms of statistical estimation, event segmentation is an unsupervised learning problem. A key claim of the model is that it can learn event dynamics while simultaneously segmenting scenes into events. In order to solve both of these problems simultaneously, we perform inference over the generative model.

As we assume that events are not directly observable, their identity and boundaries must be inferred (segmented). Given a history of scenes 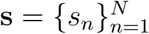, Bayes’ rule stipulates the posterior over events 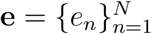 is:

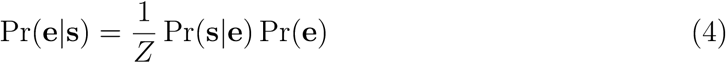

where Pr(**s**|**e**) is the likelihood of the scene history under a hypothetical event segmentation, and Pr(**e**) is the prior probability of the event sequence (i.e., the generative process for events, Eqn. 1) and *Z* is the normalizing constant. The likelihood can be decomposed according to:

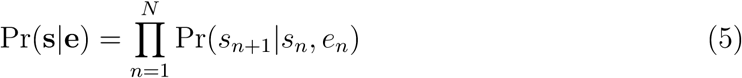

where Pr(*s_n_*_+1_|*s_n_, e_n_*) = N (**x***_n_*_+1_; *f* (**x**_1:_*_n_*; *θ_e_*), diag(*β*)) is given by Eq. 2.

As the likelihood has a Gaussian form, it can be viewed as a way of encoding (inverse) *prediction error*, which has been suggested as a key determinant of event segmentation (Reynolds et al., 2007; Zacks et al., 2011). Here, we define prediction error as the Euclidean distance between the observed and predicted outcomes. This can be thought of as a multivariate generalization of univariate prediction errors, which are typically defined as the distance between two scalar values. The Gaussian likelihood between two successor scenes (Eq. 2) is a function of the distance between the predicted scene, *f* (**x**_1:_*_n_*; *θ_e_*), and the observed successor scene, **x***_n_*_+1_. To make this point more explicit, we can express the log-likelihood for a single transition *s_n_* → *s_n_*_+1_ as follows:

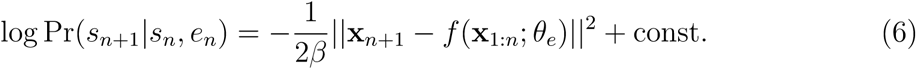

Thus, the log probability is inversely proportional to the prediction error. A low likelihood indicates a high prediction error, favoring the inference of an event boundary. Because low prediction errors correspond to high probability under the likelihood, an event model with small prediction errors will tend to be considered highly probable.^2^ It is worth noting that a prediction error can also be derived from the prior (i.e. the sticky-CRP) as well. This component of the model makes predictions as well, these predictions can be violated to varying degrees, and these violations are used to update the predictions. The posterior over segmentation considers both sources of prediction error.

Mathematically, this framing of event segmentation casts the problem as a form of clustering, in which scenes are organized into events by how well the event dynamics predict them. This is similar to prior categorization and clustering models, in which exemplars are compared to the members of previously encountered clusters and either assigned to a cluster with similar members or assigned to a new cluster (Collins & Frank, 2013; Gershman, Blei, & Niv, 2010; Love, Medin, & Gureckis, 2004; Sanborn, Griffiths, & Navarro, 2010). The key difference between these clustering models and SEM is that SEM uses event dynamics, not a set of exemplars, to determine the assignment (segmentation) of events. Because of this, two identical scenes experienced at different points in time can be assigned to different events depending on the history of scenes and events that preceded them.

In principle, event segmentation requires inference over the intractably large discrete combinatorial space of partitions. To comply with the cognitive constraint that inference is carried out online (i.e., without re-segmenting past experiences), we employ a local maximum a posteriori (MAP) approximation (Anderson, 1991; Gershman et al., 2014):

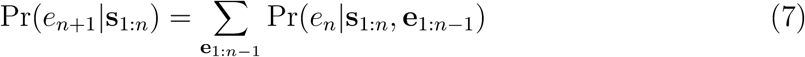

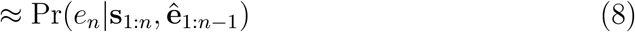

where **s**_1:_*_n_* denotes the sequence of scenes observed from time 1 to *n* and ê_1:_*_n_*_−1_ is a point estimate of the prior event segmentation defined recursively as follows:

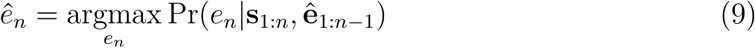

In other words, we approximate the intractable summation over **e**_1:_*_n_* with a single high probability hypothesis about the prior segmentation. For each new scene, however, we still evaluate the full posterior conditioned on this estimate. As a practical matter, this entails generating a predicted successor scene for each event in the hypothesis space and using this prediction to evaluate the likelihood of the observed scene transition. For all but the case *e_n_* = *e_n_*_+1_, this approximation assumes an event boundary (Eqn. 3), substantially reducing the computational cost.

The local-MAP approximation is commonly used in Dirichlet process mixture models of clustering (Collins & Frank, 2013; Franklin & Frank, 2019; Gershman, Monfils, Norman, & Niv, 2017; Gershman et al., 2014), and previous comparisons between the local-MAP approximation and more exact forms of approximate inference have shown them to often be highly similar (L. Wang & Dunson, 2011). It is important to note that this approximation occurs over a single forward sweep of the observed scenes, and thus does not allow for retrospective re-evaluation of the time of event boundaries. Other methods of approximate inference, such as particle smoothing, would allow for this phenomenon.

### Event Memory

We now turn our attention from the problem of event segmentation to consider the encoding of items into memory. We first define the generative process of encoding items into memory and then return to discuss how event dynamics can be used to improve memory in a reconstructive process. Consider the paired sequences of embedded scenes **x** = {**x**_1_*,..,* **x***_N_* } and events **e** = {*e*_1_*, …, e_N_* } that define the world dynamics of our generative model. Each scene vector **x***_n_* ∈ ℝ*^d^* is a real-valued vector encoding the features of the scene at time *n*, while *e_n_* ∈ ℤ is a label corresponding to the event model at the same time.

Implicitly, both **x** and **e** encode time via position because they are ordered sequences. In order to make this representation explicit, we define an unordered set of memory items *y* = {*y*_1_*, …, y_N_* } that have a one-to-one correspondence to the scene vectors **x** and events **e**. Each element *y_i_* is defined as a 3-tuple of the features of the scene, the event label, and its time index, such that *y_i_* = (**x**’*, e*’*, n*) where **x**’ is the vector of features, *e*’ is the event label and *n* is the time index. Equivalently, **x**’ = **x***_n_* and *e*’ = *e_n_* for the scene *y_i_* = (**x**’*, e*’*, n*).

We assume memory is a lossy encoding and retrieval process such that all of the components of the memory items are corrupted. Specifically, we assume an encoding process Pr(*ỹ*|*y*) creating the corrupted (encoded) memory trace 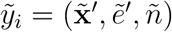 where 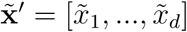, ẽ’and ñ corresponds to the corrupted memory traces of the scene features, event label, and time index, respectively. The assumption that memory traces are corrupted versions of an original stimulus is common in computational models of memory (Hemmer & Steyvers, 2009; Huttenlocher et al., 1991; Shiffrin & Steyvers, 1997) and is analogous to a capacity-limited compression (Brady, Konkle, & Alvarez, 2009; Nassar, Helmers, & Frank, 2018).

For convenience, we will assume that the corruption process for each component of each scene vector is independent, such that

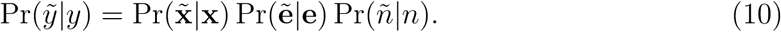

We assume Gaussian noise over the features, such that

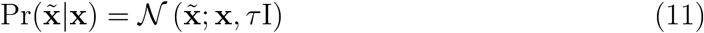

where the parameter τ corresponds to the degree of corruption noise of the feature. For event tokens, we assume that the corruption process is an asymmetric channel similar to a Z-channel (MacKay, 2003), such that the event token is either correctly encoded or that the event label is completely lost:

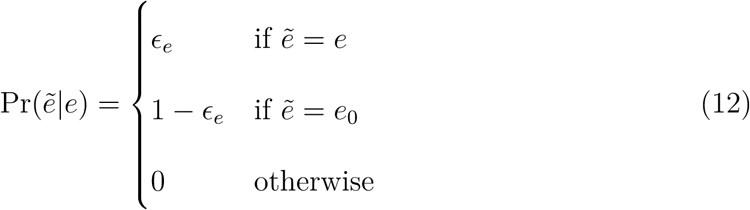

where *e*_0_ is a *null* event label corresponding to no event model (representing a loss of the event label), and where the parameter ϵ*_e_* defines the probability of retaining the event label in memory. Thus, when ϵ*_e_* = 0, the event label is completely lost from the memory trace. Corruption noise over the time index *n* is defined as a discrete uniform over the interval [*n* − *b, n* + *b*], such that

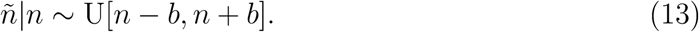

We will typically assume small values of *b*, allowing for items in memory to be flipped, but not allowing large jumps in time. Taken with the corruption of event labels, time index corruption has several consequences, including the corruption of the relative order of items within each event and the corruption of the (implicitly represented) boundary location.

We note that while we have committed to these independent corruption processes, other forms of information loss in memory are plausible, including event-label switching and non-uniform time corruption, among others. We have assumed that the corruption processes are independent of each other as a simplifying assumption, but correlations between sources of corruption noise are indeed plausible. A strong positive correlation would tend to lead to an all-or-nothing memory trace in which a memory item was strongly corrupted or not at all. Non-random forms of corruption are also possible, such as an optimal encoding process that only stores features in memory if they are not predicted by the current event model. This kind of encoding would result in the memory trace emphasizing schema-atypical features of individual scenes.

#### Reconstruction

The task of the memory model is to reconstruct *y* given a memory trace *ỹ* and the learned event dynamics. Above, we defined a generative process by which a noisy memory trace *ỹ* is created, and our model of memory is an inversion of this generative process to reconstruct the original items encoded into memory. Knowledge of the event dynamics is helpful in memory reconstruction because they are part of the generative process of the original scenes and thus contain information that can be used to aid reconstruction. This can be understood as a form of regularization: the noisy memory trace will be regularized towards the typical scenes under the event schema. A similar approach by Huttenlocher and colleagues (1991) was used to model human memory judgments of the spatial location of dots. They argued that a category prior can be used to reduce the variance of a corrupted memory trace. Here, we argue that event dynamics occupy the same role, reducing the variance of the corrupted memory trace by regularizing (that is, adding an adaptive statistical bias). This general account of event memory is compatible with other forms of corruption than those we have defined above.

To implement the memory model, we first consider the generative process of *ỹ_i_* (and its corrupted features 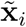 and time index *ñ*). This is defined as:

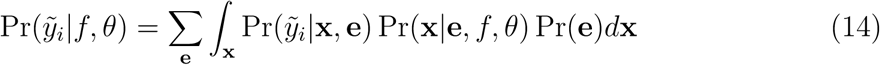

where **x** = **x**_1:_*_n_* and **e** = *e*_1:_*_n_* are the sequences of scenes and event labels, respectively. The variable θ = {*θ_e_, θ_e_*′*, …*} denotes the sets of all parameter sets that define each event’s dynamics (i.e., the weights and biases of the recurrent neural network). The three probability distributions, Pr(*ỹ_i_*|**x**, **e**), Pr(**x**|**e***, f,* θ), and Pr(**e**) correspond to the encoding process, transition dynamics and prior over events, respectively.

The goal of memory retrieval is to estimate the original scenes *ŷ* using a reconstruction process over the generative model (equation 14). Because the posterior has no closed-form expression, we employ Gibbs sampling to draw a sample of reconstructed memory traces. A complete description of our Gibbs sampling algorithm is detailed in Appendix C. At a high level, Gibbs sampling takes advantage of the conditional independence properties of the generative model, which can be seen in the graph structure (Figure 4). At each point in time, the generative process for the memory trace, the dynamics over scenes and the dynamics over events can be expressed with three functions that express the conditional independence properties (equations 10, 2, and 1, respectively). As such, we can draw samples of the memory traces, reconstructed scenes and event labels one variable at a time by conditioning on the other variables in the process. We use this sampling process as a model of reconstruction memory and do not fit it to human data.

**Figure 4.**
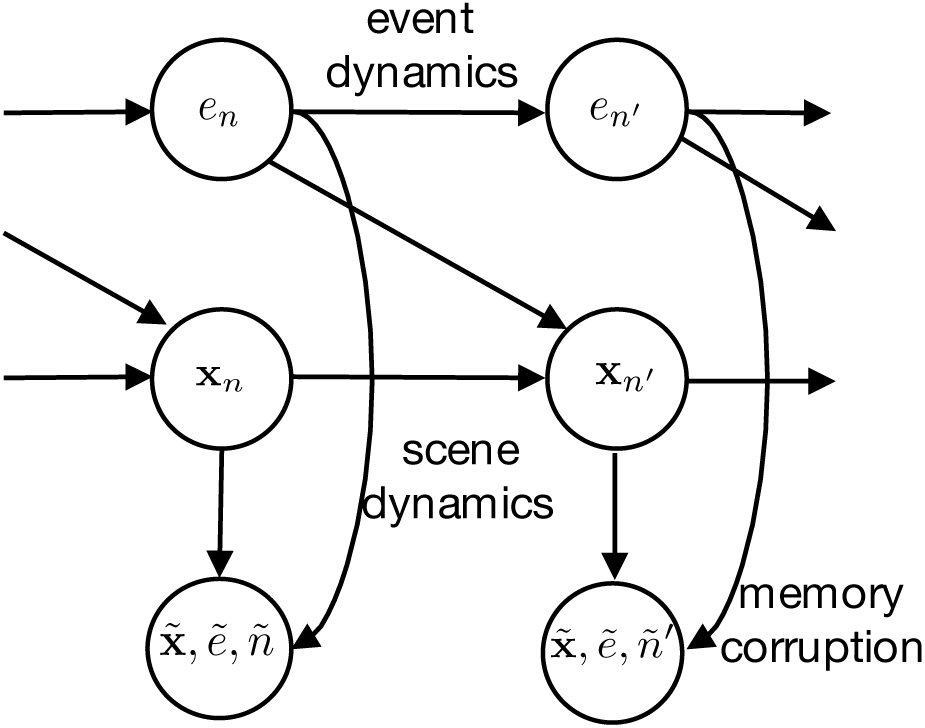
Memory corruption. A lossy memory encoding procedure stores the features of the original scene **x**, its time index *n* and event index *e* in corrupted form 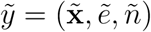. Memory reconstruction inverts this generative process to infer the original memory item.

To capture the possibility of memory failure, we augment the set of corrupted memory traces {*ỹ*} with a *null* memory *y*_0_ and define a special case of the corruption process such that

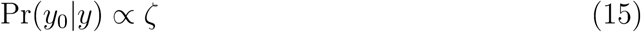

where ζ is a free parameter that controls forgetting in the model, as we discuss below. This choice of corruption process is convenient, as it has the interpretation of integration over memory items, 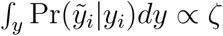 for any arbitrary value of ζ.^3^ We assume that *y*_0_ can occur multiple times in the reconstruction process, as it does not correspond to a specific memory trace, but the absence of one. Finally, we also assume that the learned event models and parameters (*f,* θ) are known because they have already been learned.

### Schematic Example

It can be helpful to walk through a toy example to provide an intuition as to how these components combine. Here, we consider a toy example with two scenes, one in which a person, Tom, asks another person, Charan, a question and a second scene in which Charan answers Tom. Symbolically, we can represent the two scenes as *Ask(Tom, Charan)* and *Answer(Charan, Tom)*. To provide these scenes to SEM, we assume that each of these two scenes is encoded using the HRR (Figure 5 B). To construct this representation, we start by constructing a representation for Tom by combining two Gaussian random vectors that represent the unique features of Tom and the property of being a person with vector addition. In the first scene, Tom is the agent and is bound to a vector that represents this role with circular convolution. This operation conserves the dimensionality of the original vectors and produces a new vector, *Tom-agent*, that represents Tom in the role of agent. We construct vectors for *Charan-patient* similarly, combining two vectors that represent the unique features for Charan and the property of being a person with vector addition, and binding the resultant vector to the patient role vector with circular convolution. Importantly, the vector that represents the “person” feature is the same for both, thus encoding similarity between the otherwise distinct symbols via vector similarity. The *Tom-agent* and *Charan-patient* vectors are combined with an *Ask-verb* vector using vector addition to create the fully embedded scene, *Ask(Tom, Charan)*. This results in a vector representation of the scene that allows us to decode each component and its corresponding symbolic role. The second scene, *Answer(Charan, Tom)* is similarly constructed.

**Figure 5.**
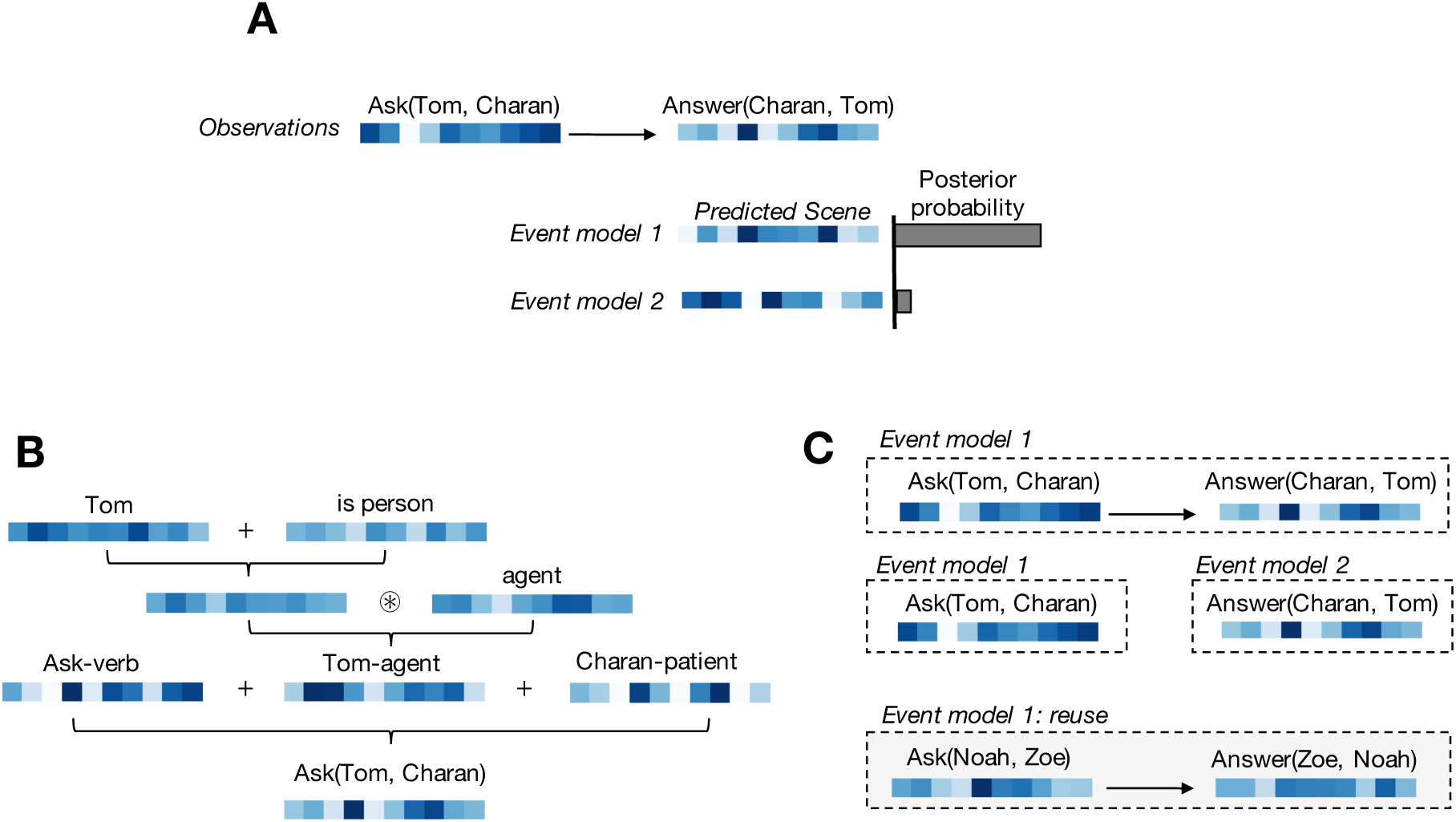
Schematic Example. *A*: The model observes a sequence of two vectors encoding the scenes *Ask(Tom, Charan)* and *Answer(Charan, Tom)*. Each event dynamics model makes a prediction for the upcoming scene, and scenes are assigned to events as a function of their similarity to the predicted scenes. *B*: Embedded scenes are constructed with a *Holographic Reduced Representation* (HRR), which combines independent features via vector addition and binds fillers to roles with circular convolution. *C*: Event clustering can assign both scenes to the same event model (top) or each scene to its own event model (middle). A learned model can be reused to predict the dynamics of a novel event token (bottom).

Given this sequence of scenes, there are two possible assignments of scenes into events. Either both scenes are assigned into one event with a learned transition between the two, or both are assigned to a unique event with one scene (Figure 5 C). How SEM makes the determination between these two assignments is determined by the likelihood of the second scene, *Answer(Charan, Tom)*, under the predictions of two possible event models (Figure 5 A; Eqn 4). In this example, SEM will deterministically assign the scene *Ask(Tom, Charan)* to one event model, and that event model will use the first scene to make a prediction about the upcoming successor scene using the dynamics learned via the recurrent neural network (Figure 3; Eqn 2). Assigning the second scene to a separate event model implies an event boundary; in this case, and SEM will use the learned initialization (*f*_0_, Eqn 3) to make a prediction for this (new) event model. Each prediction is used to calculate the likelihood of each event and combined with the sticky-CRP (Eqn 1) to calculate the posterior over event segmentations. When the predicted scene for a particular event is similar to the observed scene (i.e., low prediction error), then this will lead to a high posterior probability and the scene will tend to be assigned to that event (Figure 5 A). Conveniently, learned event dynamics can be used to explain novel events when those events are similar to previously experienced ones. For example, if the model learns the event *Ask(Tom, Charan)* followed by *Answer(Charan, Tom)*, it can generalize this pattern to the new event *Ask(Noah, Zoe)* followed by *Answer(Zoe, Noah)*, as this new event has a similar construction and event dynamics (Figure 5 C).

## Simulations

The parameter values for all of the following simulations are listed in Tables 1 and 2. The parameter values for the simulations are an implementational choice, and because the choice of parameter values interacts with the representational space of the simulations, multiple equivalent, but distinct, combinations of representational spaces and parameter sets are likely to produce the same behavior. Consequently, the parameter values only have meaning in the context of a specific representational space chosen for each task. In our simulations, the timescale of a scene, as well as the dimensionality of the representation, vary substantially. As such, we cannot make strong claims about the parameter values and their relation to cognition. Where appropriate, we have nonetheless reused parameters across multiple simulations to show that what might be interpreted as conflicting behavior falls in the same parameter space and is not driven by choosing different values for different simulations. These simulations are noted below.

The code for all simulations is available in our Github repository: https://github.com/ProjectSEM/SEM.

### Human-like segmentation on naturalistic stimuli

A key test of the model is whether it generates human-like segmentation on naturalistic tasks. Operationally, event boundaries in human studies are often defined by having subjects mark them in a naturalistic dataset, for example, while watching a video (Baldassano et al., 2017; Hanson & Hirst, 1989; Newtson & Engquist, 1976; Zacks et al., 2006, 2001). Historically, this has posed a challenge for computational models due to the difficulty of dealing with unannotated raw video data. The computational model proposed by Reynolds et al. (2007) attempted to circumvent this problem by using a carefully collected, low-dimensional motion capture dataset. Reynolds et al. (2007) evaluated their recurrent neural network model of event processing on a set of motion capture time-series, each of which contained 18 points in a 3-dimensional coordinate space measured across time with a 3Hz sampling rate. While this model was able to provide valuable theoretical insights, such as the feasibility of updating event models with prediction error, it is difficult to fully validate the model without comparing the model directly to human data.

Because of the rapid improvement in computer vision in recent years (LeCun et al., 2015), this computational issue can now be tackled; we can directly evaluate SEM on the same naturalistic dataset used in human studies. In the following simulations, we evaluate SEM on three video datasets previously used in a pair of studies by Zacks et al. (2006, 2001) to probe human event segmentation. We use the model to generate event boundaries and compare its predictions to human behavior reported in Zacks et al. (2006). We do so with fully unsupervised training: we first estimate an unstructured scene representation with a variational auto-encoder (VAE; see Appendix A for details) before providing this scene representation to the model. These unstructured scene representations are not fit to human data, but instead are used as a dimensionality reduction tool. It is important to note that the scene representation estimated by the VAE does not explicitly encode objects or bound relations between objects but can be better thought of as a dimensionality-reduction technique that takes advantage of spatial correlations in the pixel data. (And is thus simpler than the structured representational scheme presented above.) Consequently, in a video of a person making a bed, the model is not told what a person or a bed is, but has access only to what can be learned from a limited dataset. We are not claiming that people only use unstructured information while performing this task, but we nonetheless believe it is important to validate the model’s performance in an end-to-end, fully unsupervised process on naturalistic data to argue that SEM scales to realistic problems. Having said this, SEM is clearly at a disadvantage relative to human participants, who presumably have substantial prior experience with the everyday objects and actions in these videos. We will return to the role structure plays in segmentation with later simulations.

## Results

The stimulus set in Zacks et al. (2006, 2001) consists of five videos of a single person completing an everyday task, such as washing dishes, shot from a single, fixed camera angle with no edits. At a resolution of 240×320×3 pixels, each frame has more than 230,000 dimensions. We used a variational auto-encoder to generate an unstructured scene representation of 100 dimensions. We evaluated the model on each of the three videos used as stimuli in experiment 1 of Zacks et al. (2006). The model was trained on each video independently with the parameter values listed in Table 1 with 25 batches of randomly initialized weights. Figure 6 depicts a comparison between human and model boundary locations in the “washing dishes” video as well as a quantitative comparison over all three videos. Both human and model event boundaries have been binned in 1-second intervals, as was reported in Zacks et al. (2006). Qualitatively, there is good agreement between the model’s maximum *a posteriori* (MAP) estimates of boundaries and the population of human subjects, with several of the major peaks in the group data corresponding to a model boundary (Figure 6, Top). For example, the most commonly marked boundary for human subjects in the “washing dishes” video occurs at 34 seconds, when the actor in the video approaches the sink. At this time-point, both 43.8% of humans and 44% of model batches mark a boundary. Both frequently note a boundary at 131 seconds (humans boundary frequency: 29.2%, model boundary frequency: 48%), when the actor opens the dishwasher. Notably, both of these changes correspond to larger changes in the visual scene. The model tends to miss more subtle changes that rely on prior knowledge. For example, the human-rated boundary at 79 seconds during which the actor turns on the sink (boundary frequency: 27.1%), is missed by the model (boundary frequency: 0%).

**Figure 6.**
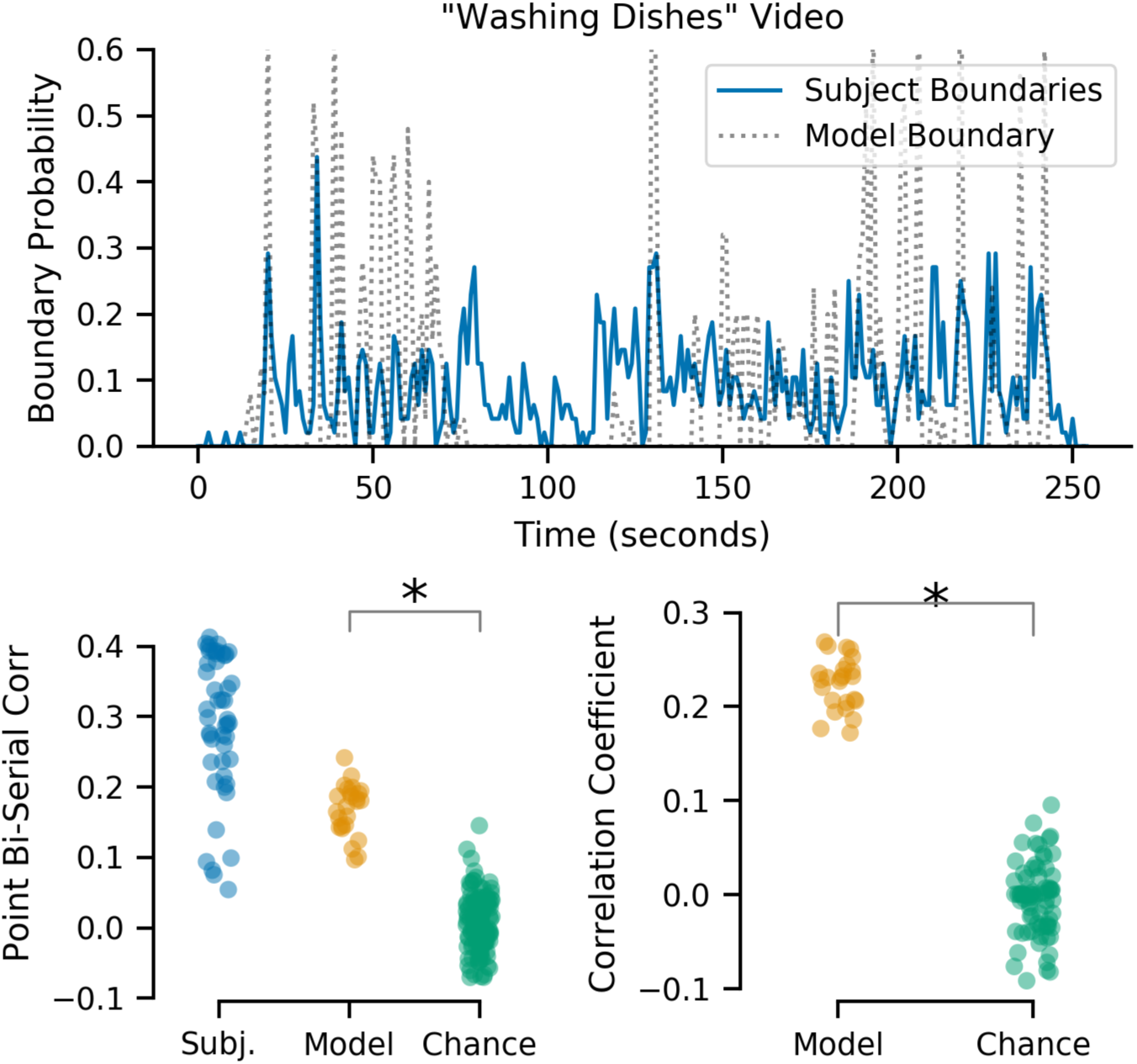
Video Segmentation. *Top*: SEM generates human-like boundaries. The model’s MAP estimated boundaries are shown for the “washing dishes” video averaged over all batches and compared to human segmentation frequency. *Bottom Left*: Point biserial correlation coefficient for human subjects (blue), the model (orange) and a permutation distribution (green), comparing discrete boundaries to aggregated human data. Model segmentation falls within the range of human performance and is above that expected by chance. *Bottom Right*: Correlation coefficient between model boundary log-probability and human-rated boundaries (orange) compared to correlations generated by permutation testing (green). For clarity, only 5 of the 1000 permutations are shown.

We assessed the model predictions quantitatively using the point-biserial correlation between the model’s MAP boundary estimates and the grouped subject data. Point-biserial correlation is a similar metric to the one used by Zacks et al. (2006) to assess the segmentation of each subject. In this naturalistic dataset, human subjects had an average point-biserial correlation of *r_pb_* = 0.29 (SEM=0.014), reflecting both a meaningful relationship between rated boundaries across the population as well as substantial between subject variability. The point-biserial correlation of the model was *r_pb_* = 0.168 (SEM=0.013), which places the model below the average value of human segmentation but within the middle 75% of the distribution of human ratings (Figure 6, Bottom Left; middle 75% of observations: [0.139, 0.400]). Thus, by falling within the same range of observed values, the model’s performance is comparable to that of human subjects. We further assessed how likely this result would occur by chance by means of permutation testing: we permuted the order of the MAP boundaries of each batch 1000 times and calculated the point-biserial correlation for the permuted samples to create a chance (null) distribution. The model’s average point-biserial correlation of 0.168 was larger than would be expected by chance (95% CI of null permutation distribution: [-0.067, 0.080]).

As the model produces a boundary probability at each time step, we can make a more sensitive comparison between model and human behavior by correlating the model’s log-probability of an event boundary with human boundary frequency. Doing so, we find a correlation coefficient of *r* = 0.22 (SEM=0.017), which we evaluated for statistical significance via permutation testing (again, creating 1000 permutations and calculating the correlation coefficient of the permuted data) and find that the mean correlation coefficient of *r* = 0.22 was well above what we would expect by chance (95% CI of null permutation distribution: [-0.071, 0.074]; Figure 6, Right). Thus, these two quantitative measures suggest that the model’s segmentation captures meaningful variance in the human data.

### Event boundaries and community structure

A natural pair of questions to ask is how SEM determines an event boundary and what features in the environment influence segmentation. In the task simulated above, this is difficult to ascertain as the environment was not directly manipulated to probe these questions. Thus, while we demonstrated that the model and human participants delineate similar boundaries in a naturalistic task, we are not able to draw strong conclusions about which statistical features of the dataset precipitate segmentation. We therefore turn our attention to controlled experiments to characterize segmentation in SEM and examine its determinants.

One such determinant that has previously been identified is prediction error (Reynolds et al., 2007). Humans show a decrease in predictive accuracy across an event boundary in naturalistic tasks (Zacks et al., 2011) and generally respect statistical structure (Avrahami & Kareev, 1994; Baldwin et al., 2008). Moreover, surprising occurrences in human tasks influence event perception (Newtson, 1973), suggesting that the predictability of scenes is important for segmentation. Given the formulation of event segmentation in our model as probabilistic inference (Equation 4), the property that surprising or unpredicted stimuli will produce event boundaries is axiomatic. The model is sensitive to both prediction error and the uncertainty of prediction, a property that can be derived analytically (see Equation 6).

Less apparent, however, is how the model responds to other types of statistical structure. At first glance, the role of predictive inference in event segmentation might suggest that humans rely strictly on surprising outcomes to drive segmentation. However, Schapiro et al. (2013) identified community structure as a feature that drives human event segmentation, even when controlling for predictability. In this context, a community refers to a group of nodes in a graph that is densely interconnected. Schapiro et al. (2013) showed subjects a sequence of stimuli drawn from a graph that had multiple communities, and subjects were asked to note transition points between stimuli. An important feature of the task is that transitions were equated for probability, such that a community transition was no more or less likely than any other individual transition. This was done to rule out the possibility that subjects rely solely on unpredictability in the environment to identify event boundaries.

Nonetheless, subjects preferentially marked event boundaries at community transition points, respecting the graph structure. Schapiro and colleagues further demonstrated that a recurrent neural network trained on the same sequence of stimuli developed internal representations that were similar for community members. Within the network, representational similarity between sequential items decreases at event boundaries, potentially acting as a signal that a boundary has occurred.

Like the Schapiro model, SEM is also sensitive to community structure. To demonstrate this, we simulated the model on 1400 stimuli generated by a random walk on the graph used in Schapiro et al. (2013) in a sequence exposure phase, followed by 600 stimuli from randomly drawn Hamiltonian paths in a parsing phase.^4^ Consistent with the previously reported human behavior and the Schapiro model, SEM has higher boundary probability for a community transition than a non-community transition, across both the training trials and in the Hamiltonian paths of the parsing phase (Figure 7b).

**Figure 7.**
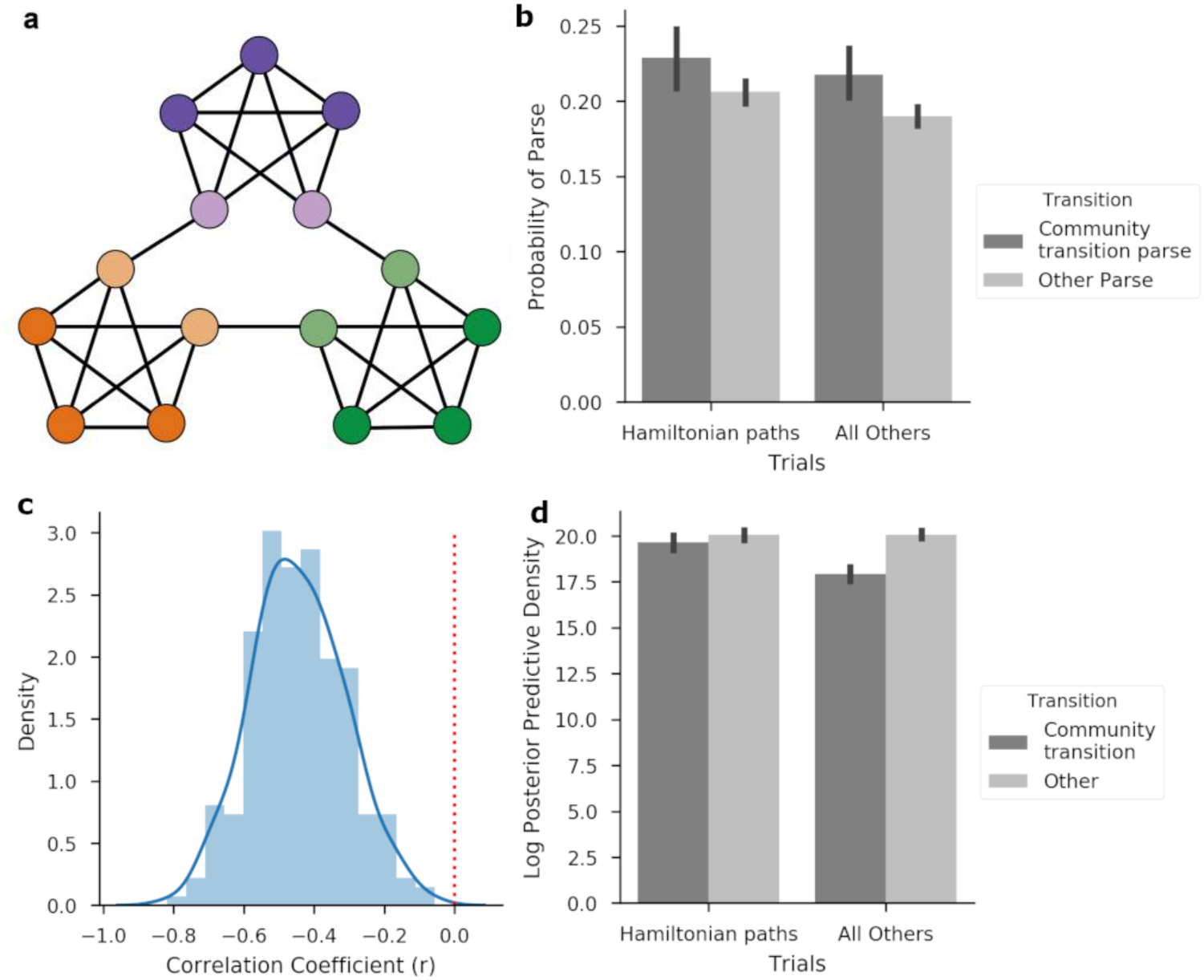
Event boundaries at graph community boundaries. (a) The graph structure of the transition matrix, adapted from (Schapiro et al., 2013). Sequences of stimuli were generated by drawing from walks through this graph. (b) The probability of a parse (event boundary) is shown between pairs of items that reflect a community transition (dark grey) or within a community (light grey) for Hamiltonian paths through the graph (left) and all other trials (right). (c) Pearson’s r coefficient between the log posterior predictive density of each scene and the boundary probability is shown for the sample of simulations. (d) The log posterior predictive density for each scene. Lower values correspond to greater surprise under the model.

It is important to note that boundary probability and prediction error are related in SEM, such that more surprising scenes are more likely to lead to an event boundary (Figure 7c). Consequently, the average log probability of each successive scene, a Bayesian measure of prediction error,^5^ is lower at community transitions than at other transitions, meaning that community transitions are more *surprising* than non-community transitions (Figure 7d). This might be seen as surprising giving the equating of transition probability in the task. However, predictability in the environment is not the same as predictability from the point of view of an agent, and SEM’s generative model is not equivalent to the generative process of the task. Furthermore, the recurrent neural network model proposed by Schapiro and colleagues relies on similar computational principles as the recurrent neural network we used to model event dynamics. Thus, the two should be sensitive to similar features of the task. In short, this simulation reconciles the apparently conflicting claims that segmentation can be driven by prediction error (Reynolds et al., 2007) and that prediction can be driven by community structure even when objective predictability is equated (Schapiro, Turk-Browne, Norman, & Botvinick, 2016): even after equating the objective predictability for an all-knowing agent, an agent who has learned the structure of the environment using an approximate model may well experience spikes in prediction error at community transitions.

### Generalizing structure

Thus far, we have shown the model can segment data in line with human judgments, and does so in a way that reflects graph community structure. However, this should be an unsurprising result: simpler computational models either relying on recurrent neural networks alone (Reynolds et al., 2007; Schapiro et al., 2013) or latent state inference alone (Baldassano et al., 2017; Goldwater et al., 2009) can produce human-like segmentation in multiple domains. This is because both recurrent neural networks and latent state inference encode temporal structure, and while they represent this structure differently, for segmentation alone either is sufficient. This raises the natural question of why is it valuable to encode multiple different types of structure and under what conditions would we expect to see the benefits of that structure?

A key insight is that temporal clustering (i.e., grouping scenes into events) and dynamical systems (i.e., the event dynamics) represent different forms of temporal structure. Clustering models group similar observations together, allowing them to pool information efficiently and generalize rapidly (Collins & Frank, 2013; Gershman et al., 2010, 2014; Sanborn et al., 2010), and this data-efficiency can be significant in sequential problems (Franklin & Frank, 2018) like those that define event cognition. However, clustering scenes into events alone does not represent order information within individual events. Were we to model events as an unordered collection of scenes, our event model would be unable to differentiate between the beginning and end of an event, nor would it be able to differentiate between an event and the same event experienced backward in time.

The previously proposed recurrent neural network models of event segmentation (Reynolds et al., 2007; Schapiro et al., 2013) have a complementary set of benefits and limitations. These models do learn the sequential transition structure of individual events and represent event boundaries within their representations of this structure. The demarcation of event boundaries in these models is implicit and these models do not have a mechanism to encourage a data-efficient pooling of observations. Moreover, when trained continually, these models tend to suffer from catastrophic interference, or the tendency of newer tasks to interfere with the knowledge of older tasks (Goodfellow, Mirza, Xiao, Courville, & Bengio, 2013; McCloskey & Cohen, 1989). Conditioning learned event dynamics on an event label presents a potential solution to this. SEM represents both of these forms of temporal structure. Combined with the third form of structure that we have not discussed thus far—the logical structure of individual scenes encoded in the vector space embeddings—the model is capable of sophisticated generalization.

To demonstrate this, we present here a simple example: an event structure that is defined by (1) a person asking a second person a question, followed by (2) the second person responding to the first person. Symbolically, we can represent this event as *Ask*(*Tom, Charan*) → *Answer*(*Charan, Tom*), where we have named the two people Tom and Charan and the expression means that Tom asks the question and Charan answers (Figure 8).^6^ SEM can learn this structure and generalize it to new fillers (i.e., other people) that it has not encountered. Specifically, the model is sensitive to the structural form of the event, without regard to role/filler assignments, and generalizes this structural form to new events. This depends both on the structured vector representations of scenes as well as its internal estimate of the event dynamics. Critically, SEM can also use this structure to detect when there is a change and assign observations to a new event.

**Figure 8.**
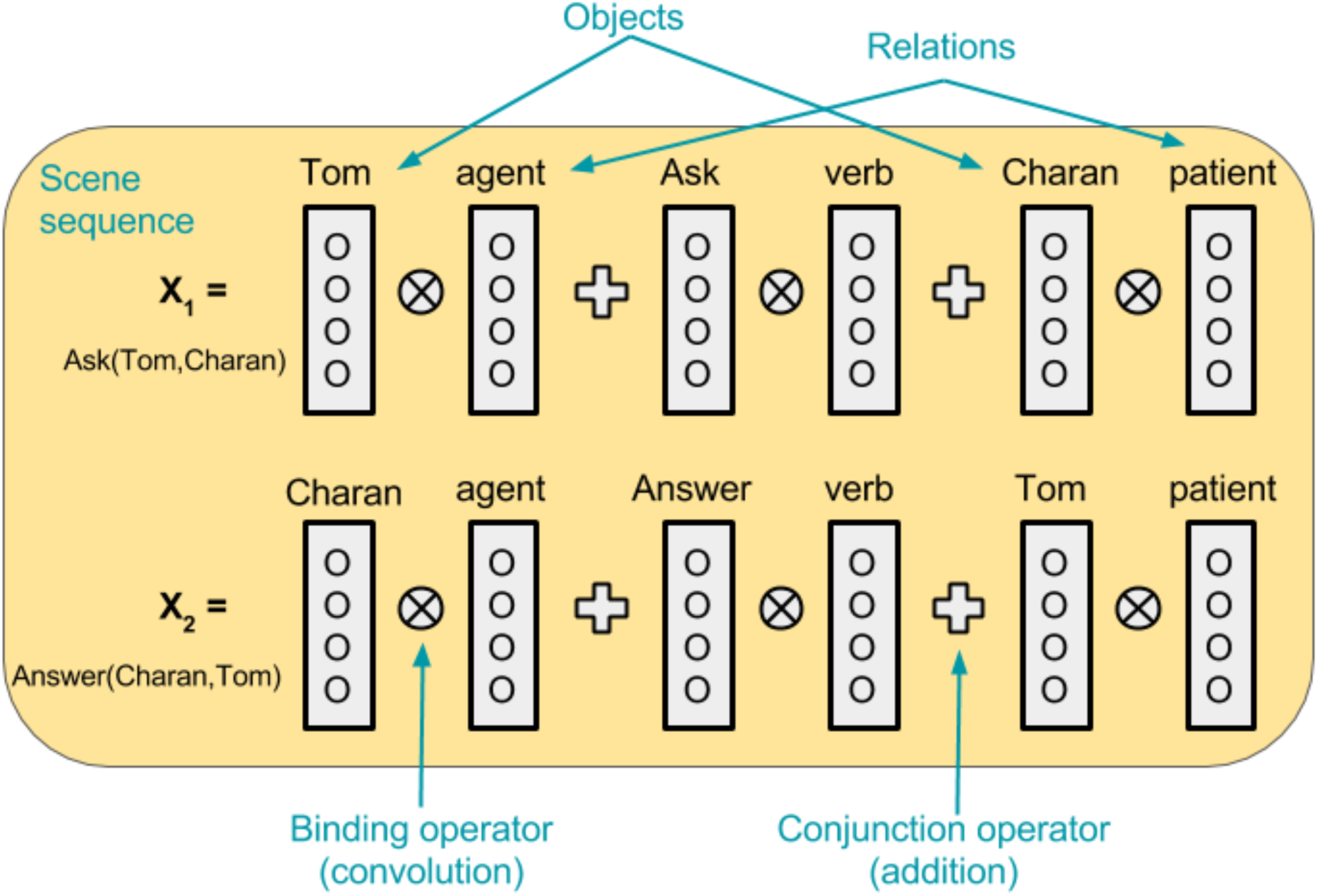
HRR of the question/answer task. Each filler is bound to its role by circular convolution. Role/filler bindings are composed of with vector addition to produce scene vectors.

We examined the probability of an event boundary between *Ask*(*A, B*) and *Answer*(*B, A*) for arbitrary fillers of *A* and *B*. Here, we are concerned with the use of structure, and therefore have pre-trained SEM with examples of the sequence *Ask*(*A, B*), *Answer*(*B, A*). This simulates event structures that are already known. SEM was pre-trained on 3 unique sequences of *Ask*(*A, B*), *Answer*(*B, A*) with 6 unique fillers. Each scene vector was composed using the HRR as previously described. Each independent feature was represented with a spherical, zero-mean, Gaussian random vector (**x** ∼ N (0*, d*^−2^I)) and similarity between fillers was encoded with a shared component. For example, the vector representing the symbol *Ask* is encoded by the vector addition of a common feature to all verb-symbols (itself a random vector) and a unique random vector to the symbol. Due to the simplicity of the event dynamics, we estimated the event dynamics without recursion and replaced the GRU layer in our function approximator with a non-recurrent, but otherwise equivalent, layer.

We simulated SEM on five test sequences, probing the event boundary probability between the first and second scenes. We also probe the probability that the second scene belongs to a new event model (as opposed to a new instance of the previously experienced event). These metrics indicate how the model partitions scenes into events and are a proxy for generalization. The first sequence, *Ask*(*Tom, Charan*) → *Answer*(*Charan, Tom*), was included in the pre-training sequences and acts as a negative control (i.e., low probability of an event boundary and a new event). It is worth noting that both *Tom* and *Charan* have different relational roles in the first and second scenes. If SEM were only sensitive to the change in relational roles, then this would precipitate an event boundary. However, as SEM learns transitions between structured scenes, this change is unproblematic as it is predictable.

A second sequence, *Ask*(*Tom, Charan*) → *Chase*(*Dog, Cat*), deviates from the event structure and acts as a positive control (i.e., high probability of an event boundary and a new event model). As expected, SEM assigns a low boundary probability to the probe *Answer*(*Charan, Tom*) (Fig 9) and a high probability of both an event boundary and a new event model for the probe *Chase*(*Dog, Cat*).

**Figure 9.**
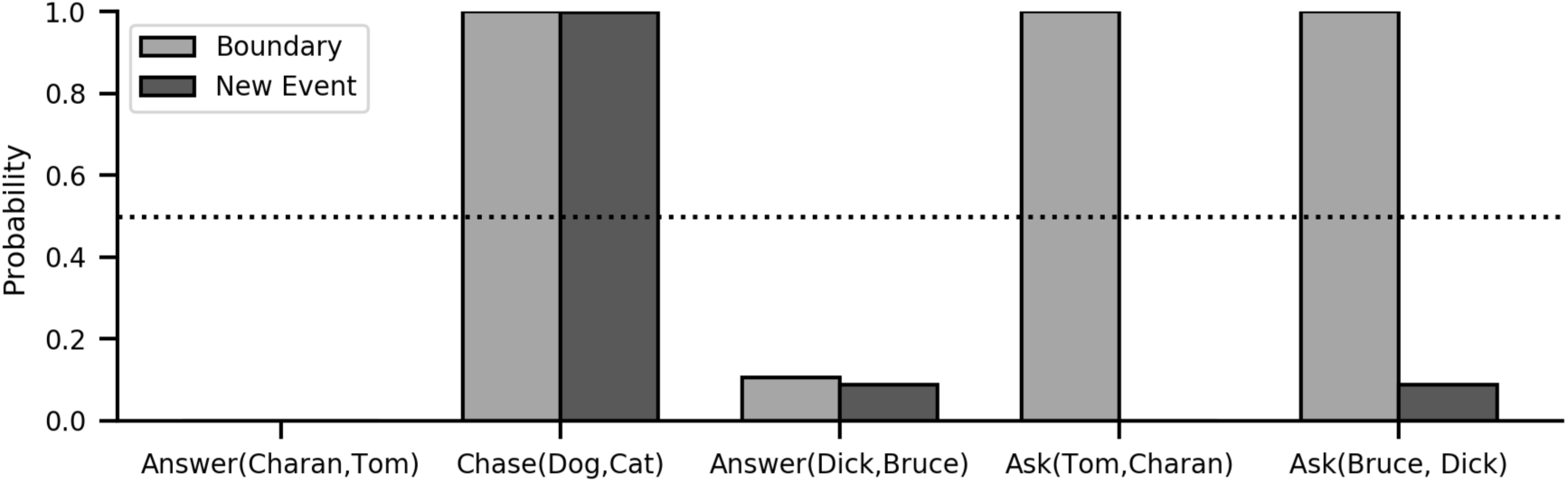
Generalization task. The probability assigned by the model for an event boundary (light grey) and a new event model (dark grey) are shown for four test events. Each event begins with the scene *Ask*(*Tom, Charan*) and ends with the denoted scene, except for the event *Ask*(*Bruce, Dick*) → *Answer*(*Dick, Bruce*). From left to right, the probabilities are shown for an event in the training set, a structurally dissimilar event, a structurally similar event with novel role/filler bindings, a control where the original event was restarted, and the beginning of a structurally similar event with novel role/filler bindings.

A key demonstration is generalizing the learned structure to arbitrary fillers. To show this, we provided SEM with the sequence *Ask*(*Dick, Bruce*) → *Answer*(*Bruce, Dick*), which used fillers (*Dick* and *Bruce*) that were held out of the pre-training set for this purpose. Here, SEM assigns a low, but non-zero, probability both that an event boundary has occurred (18.0%) and that the probe belongs to a new event model (12.2%). Equivalently, this corresponds to a high probability that both scenes are assigned to the correct event with the correct model (82.0%), reflecting a generalization of the relational structure. This highlights that SEM is sensitive to both the structured and non-structured features of the scene. Non-structural features are novel, and the probability of the previously learned event under the model is relatively lower. Consequently, SEM makes the prediction that people rely on both structural and non-structural features for segmentation. Nonetheless, we would expect the structured features to dominate for highly familiar event types as this structure of the event is thought to be more predictable across time (Richmond & Zacks, 2017).

Next, we look at how SEM reuses events multiple times. SEM uses Bayesian inference to identify events and supports re-using an event model following an event boundary. To illustrate this, we provide SEM with the sequence *Ask*(*Tom, Charan*) → *Ask*(*Tom, Charan*), in which the first item is repeated. Here, SEM infers an event boundary (even as the two vectors are identical) but does not assign the second item to a new event. Instead, SEM infers that the original event has been restarted. Interestingly, this is a property of the structure of the event, and SEM will reuse an event following a boundary based on this structure. This can be shown by providing SEM with the sequence *Ask*(*Tom, Charan*) → *Ask*(*Bruce, Dick*), which reuses the event with a pair of novel fillers. SEM infers an event boundary between the two items and re-uses the previous event (i.e., infers a low probability of a new event model), again showing structure sensitivity. Thus, by inferring the event type in addition to learning the dynamics over sequential scenes, SEM can rapidly re-use event dynamics in novel events with a shared relational structure.

This reuse of previous event models also highlights how SEM uses predictions of future scenes, and not changes in the vector from scene to scene. To demonstrate this, we also simulated a reduced variant of SEM in which the event dynamics are simulated as a stationary process, similar to a hidden Markov model (Rabiner, 1989). We repeated the simulations of our negative (*Ask*(*Tom, Charan*) → *Answer*(*Charan, Tom*)) and positive controls (*Ask*(*Tom, Charan*) → *Chase*(*Dog, Cat*)) with this reduced model. As can be seen in Figure 10, the model does not strongly differentiate between the two cases. For the negative control, the model infers the probability of an event boundary at approximately chance (0.5) and infers a low probability that the second scene is from a new event model. This occurs because the model does not contain order information and thus cannot differentiate between a new event model and a new scene from the same event. For the positive control, both the boundary probability and the probability of a new event model are higher for the scene *Chase*(*Dog, Cat*) than for *Answer*(*Charan, Tom*), reflecting the dissimilarity between the scenes; however, the lesioned model is worse than the intact model at inferring that *Chase*(*Dog, Cat*) belongs to a new event, as the lesioned model does not learn the appropriate structure.

**Figure 10.**
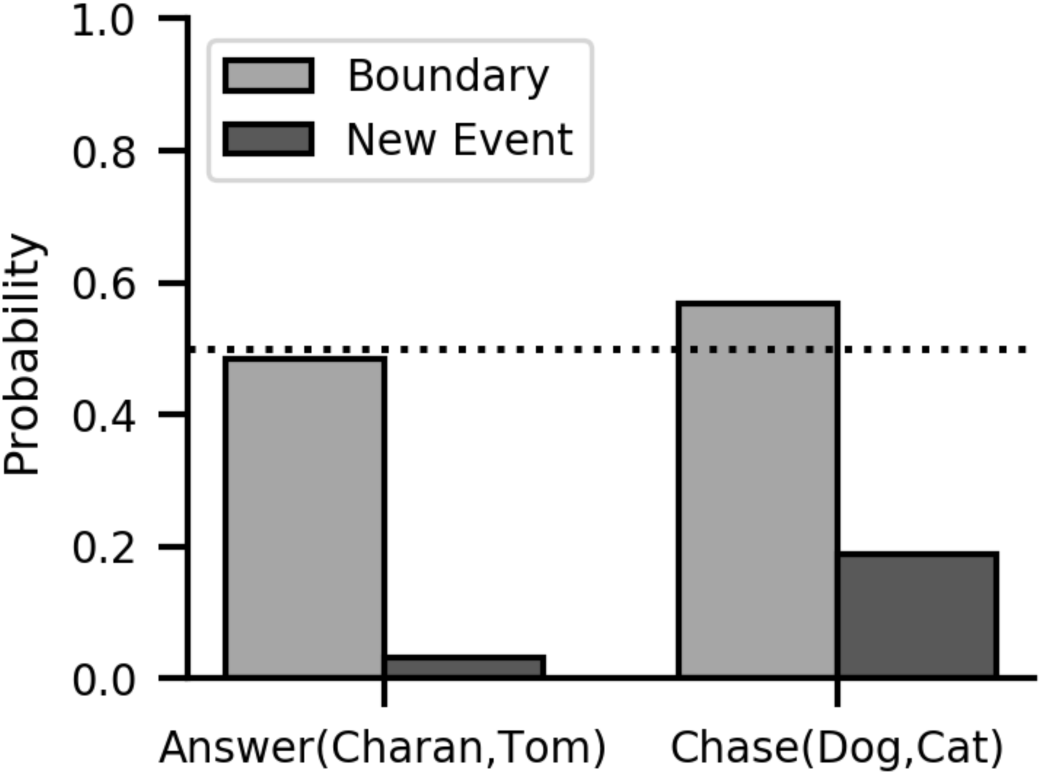
Lesioned event dynamics. SEM was lesioned such that it does not learn the dynamics but only a stationary distribution over scenes. The figure shows the probability assigned by the lesioned model for an event boundary (light grey) and a new event model (dark grey) for two test events. Both events begin with the scene *Ask*(*Tom, Charan*) and end with the labeled scene. From left to right, the probabilities are shown for an event in the training set and a structurally dissimilar event. Lesioning the model impedes its ability to see *Answer*(*Charan, Tom*) as part of the same event as *Ask*(*Tom, Charan*), and it also impedes its ability to infer a new event model for *Chase*(*Dog, Cat*) (see text for explanation).

Event segmentation plays a further role in the structured generalization by efficiently pooling similar event instances together and separating them from irrelevant information. This has an important protective function, as SEM is constantly learning event dynamics. To demonstrate this, we compare SEM to a second reduced model that does not differentiate between events or delineate event boundaries,^7^ and examine the prediction error of the model in a well-known event following a novel, competing event. The model was pre-trained as before and then shown the event *See*(*Dog, Cat*) → *Chase*(*Dog, Cat*) before being presented with the previously experienced event *Ask*(*Tom, Charan*) → *Answer*(*Charan, Tom*). We then measured the prediction error of the second scene in the event *Answer*(*Charan, Tom*), normalized between zero and one. SEM predicts the scene accurately, with a normalized prediction error of 0.037 (Figure 11). In contrast, the reduced model does substantially worse and has a normalized prediction error of 0.97. As SEM and the reduced model learn event dynamics with the same function and have seen the experienced the same sequence of scenes, the partitioning of data afforded by event clustering allows SEM to more efficiently learn and reduce interference.

**Figure 11.**
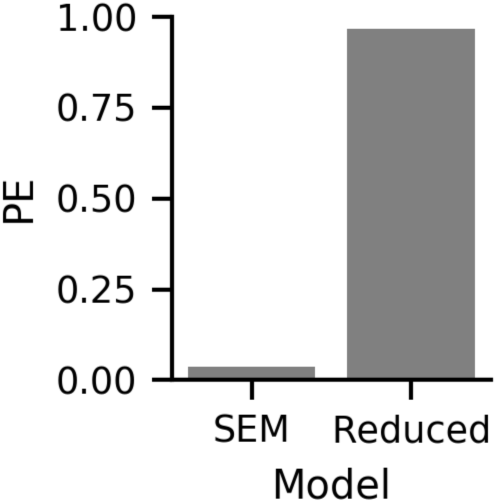
Lesioned event segmentation. Results from the intact SEM model and a reduced model without event segmentation. The figure shows normalized prediction error when the model is presented with the previously experienced event *Ask*(*Tom, Charan*) → *Answer*(*Charan, Tom*), following the presentation of a distractor event. Prediction error in response to *Answer*(*Charan, Tom*) is substantially higher in the lesioned model (see text for explanation).

Taken together, these simulations paint a picture in which the model learns the temporal and relational structure of individual events and uses this both to determine event boundaries and to generalize to novel, but structurally similar, events. Empirically, we would expect people to pool events with the same relational structure, both representing them more similarly and pooling their learned transition structure. We would also expect subjects to delineate the same boundaries for a sequence of events with a learned relational structure but novel fillers.

### Structured memory inference

We now turn to the memory predictions of the model. We first focus on false memories, which have long been thought to be a consequence of reconstructive memory (Bartlett, 1932; Roediger & McDermott, 1995), and are a natural prediction of the reconstructive memory model.

In a classic finding, Bower et al. (1979) found that subjects who read multiple similar stories drawn from the same *script* would falsely recall portions of the story that weren’t present in the original story. For example, subjects might read a story in which a character *John* goes to the doctor and reads a magazine before seeing the doctor, and a second, similar story in which a character *Bill* goes to the dentist and has to wait. Given these two stories, which shared a common event structure and were drawn from the same script, a subject might falsely recall the detail that *John* has to wait to see the doctor, a likely inference that was nonetheless not stated in the original story. This tendency to recall unstated memory items increased with the number of stories subjects read from the same script, suggesting that subjects were recombining elements of stories with a shared event representation in the recall process. This process may be adaptive to the degree that it reflects inferences about unexperienced scenes that nonetheless occurred.

We use the reconstructive memory model to generate structured false memories with a paradigm similar to the one presented in Bower et al. (1979). In the original stimulus set, stories with the same script had highly similar beginnings and endings and had a combination of structurally similar sentences and distinct, story-specific sentences in the middle that suggested a similar event trajectory. To model this, we defined a simplified set of five stories from three different scripts, each of which was comprised of four scenes (Table 4). Three stories belonged to the same script and shared a common structure but used different fillers within the structure. These sentences all shared the first and last scene, and any two of the three shared a third scene.

**Table 3.** Stimuli for generalization task. For each of the five test cases, the probability of an event boundary was measured between the 1st and 2nd scenes.

| Test Case | 1st Scene | 2nd Scene |
| --- | --- | --- |
| Training example | Ask(Tom, Charan) | Answer(Charan, Tom) |
| Structure violation | Ask(Tom, Charan) | Chase(Dog, Cat) |
| Novel role/filler binding | Ask(Bruce, Dick) | Answer(Dick, Bruce) |
| Structure Re-use | Ask(Tom, Charan) | Ask(Tom, Charan) |
| Structure Re-use with novel fillers | Ask(Tom, Charan) | Ask(Bruce, Dick) |

**Table 4.** Stimuli for Bower task. Each row denotes a story with comprised of four scenes, representing a simplified narrative containing relational structure. Each of the stories in script 1 contain common structural elements, with the same beginning and endings and a shared 2nd or 3rd scene, but with different fillers in the roles.

| Script | 1st Scene | 2nd Scene | 3rd Scene | 4th Scene |
| --- | --- | --- | --- | --- |
| 1 | Goto(John, Doctor) | Checkin(John) | Read(John, Magazine) | Treat(Doctor, John) |
|  | Goto(Bill, Dentist) | Read(Bill, Magazine) | Wait(Bill) | Treat(Dentist, Bill) |
|  | Goto(Natasha, Chiropractor) | Checkin(Natasha) | Wait(Natasha) | Treat(Chiropractor, Natasha) |
| 2 | BuyTicket(Jill, Movie) | FindSeat(Jill) | Buy(Jill, Popcorn) | Watch(Jill, Movie) |
| 3 | Wakeup(Sarah) | GetDressed(Sarah) | Eat(Sarah, Toast) | Leave(Sarah) |

The manipulation of interest is whether providing SEM with multiple similar stories will increase the probability of a script producing semantically valid false memories under the reconstruction model. Concretely, we test for the reconstructive memory probability of the probe cue *Wait(John)*, a scene that does not incur in any story, but is syntactically correct and consistent with the *Goto(John, Doctor)* story in script 1 (Table 4).

We provided SEM with three stories, including either one, two or three stories from script one. In our simulations, we used the HRR as previously described to encode each scene but provided no pretraining or other semantic knowledge to the model. Each verb (e.g., *Goto*) is a random Gaussian vector that can be decomposed into two vectors, one corresponding to the unique features of the verb and another corresponding to its structural role. For example, the vector embeddings of *GoTo* and *Read* are each the sum of a shared feature vector corresponding to the structural role as a verb and a unique feature vector that distinguishes them from other tokens. Consequently, all verb vectors are similar (close in vector space) while nonetheless distinct due to their unique features. Agents (e.g., *Natasha*) and objects (e.g., *Popcorn*) are encoded similarly. SEM was first provided the scenes for each story to learn the event dynamics and infer the event labels. Because we are not examining segmentation with these simulations, all of the scenes from a single story were assumed to belong to the same event and SEM was tasked with inferring a single label for them. It is worth noting the segmentation problem present in Bower et al. (1979) is trivial: stories were presented with clear external cues for the beginning and ends with intervening time between each story.

We then used the Gibbs sampling algorithm as previously described (see Table 1 for parameter values) to simulate reconstructive memory. We modeled a two-alternative forced-choice recognition memory test, comparing the script-consistent false memory probe *Wait(John)* with a syntactically valid but script-inconsistent memory probe *GetDressed(John)*. This memory score can be described as the expectation of the comparison

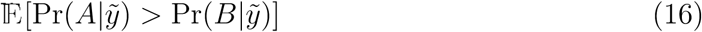

under the memory model, where *A* and *B* are memory probes. Let *f* (**x**) = Pr(**x**|*ỹ*) be the recognition memory probability under the model, and the expectation above defined as an expectation over *f*. Our reconstruction samples 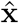 (equation 18) are drawn from samples of *f*. Thus, if we assume a function 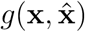 to be monotonically related to a sample of *f*, we can evaluate the ordinal comparison of the two memory probes on *g* and approximate the memory score with the average of *N* samples:

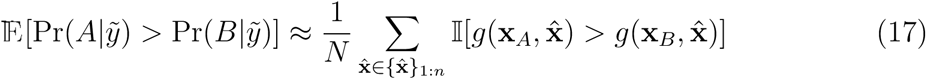

where 𝕀[·] = 1 when its argument is true, and 0 otherwise. Here, we choose 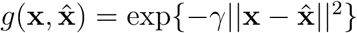, with γ = 2.5.

Figure 12 shows the results of our simulations. The script-consistent false memory probe *Wait(John)* was compared to the script-inconsistent probe *GetDressed(John)*. The recognition memory probability for the script-consistent probe increased monotonically with the number of script instances in the stimuli, similar to the behavioral finding of Bower et al. (1979). For a single script instance, this was below chance, as the comparison probe *GetDressed(John)* was more reflective of the training set than the script consistent probe. Additionally, the reconstruction process improved as a function of the number of script instances. This can be seen in the frequency with which the original memory traces were included in the reconstruction sample, which increased monotonically with the number of script instances (1 script instance: 0.82, 2 script instances: 0.88, 3 script instances: 0.92). Under SEM, these two effects are related due to the regularizing effects of the reconstructive process.

**Figure 12.**
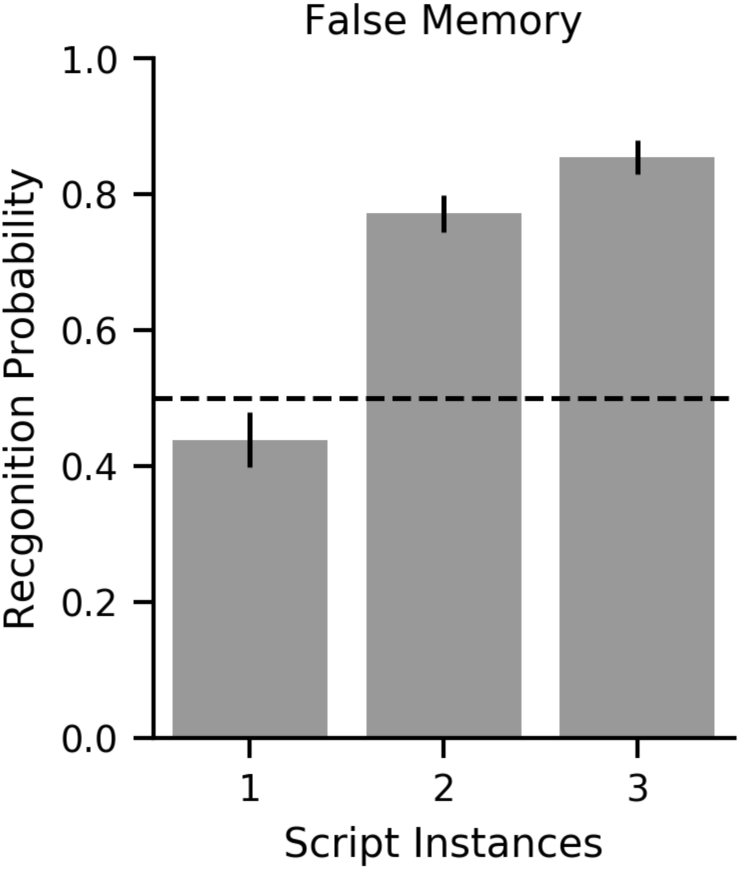
False memory simulations. The recognition memory probability for a script-consistent false memory probe is shown as a function of the number of stories of the same *script* during learning. Chance (50%) is denoted by a dotted black line.

### Event boundaries and working memory

An empirical consequence of event boundaries is that items that occur in an ongoing event are better remembered than items that occur immediately before an event boundary (Pettijohn & Radvansky, 2016; Radvansky & Copeland, 2006; Radvansky et al., 2011; Radvansky, Tamplin, & Krawietz, 2010). A study by Radvansky and Copeland (2006) had subjects remember items in a virtual environment as they moved from room to room. In each room, subjects put an item they were carrying (but was not visible to them) on a table and picked up a second object, before carrying it to another room. Subjects were given a memory probe either immediately after walking through a door (*shift condition*) or at an equidistant point in a larger room (*no-shift condition*). Overall, subjects remembered items better in the no-shift condition than in the shift condition, suggesting that the act of walking through a door interfered with the item memory. This appears to be distinct from context-change effects that have been observed in long-term memory (e.g., Godden & Baddeley, 1975), because returning to the original room in the shift condition did not eliminate the memory decrement (Radvansky et al., 2011).

According to the Event Horizon Model (Radvansky, 2012), subjects are less able to answer the memory probe in the shift condition because working memory has privileged access to the ongoing event model, and walking through the door triggers an event boundary. Why would working memory have privileged access to the ongoing event model? From the perspective of SEM, event models are used to predict upcoming scenes, and maintaining an active representation of the most recently used event model constitutes caching in memory the results of an expensive inference process. Put more simply, in order to use an event model to make predictions, a person has to know which model to use. This persistent representation is a source of information that a normative agent will use in any computational task where it is relevant, including memory reconstruction.

In SEM, there is no representation of the active event model during reconstructive memory. However, we can simulate these effects by directly providing the model with additional information about the identity of the ongoing event, which we simulate by increasing the precision of the corrupted event label 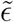. Specifically, we increased the event label precision parameter ϵ*_e_* from 0.25 to 0.75 for ongoing events. This precision parameter is only used in the creation of the corrupted memory trace and is not used during reconstruction, and this change in memory precision acts as an input to the model. While we have chosen to model the active event model with this change in memory precision, our key claim is that the active event provides a source of information to reconstruction. Other mechanistic implementations, such as an informative initialization point in reconstruction or as an external, separately defined input that biases reconstruction, would also be consistent with this account.

We simulated this hypothesis using a simplified variant of the Radvansky and Copeland (2006) task. In the original task, subjects navigated from room to room in a virtual environment. We simplified this to a stereotyped set of scenes, using a similar sequence of structured scenes as in the previous memory experiment and in the generalization simulations. To model the interactions in each room, we provided SEM with observations of (1) entering the room, (2) putting an object down (3) picking up the next object and (4) leaving/crossing the room. Each of these scenes is composed of a verb (e.g., *Enter*) and context that corresponds to an individual room (Table 5). We assumed that the object to be picked up was observed in the first two scenes. For balance, we assumed that the object that was put down was observed in the second two scenes. We further assumed that the pickup object was bound to the verb *Pickup* in the third scene. Each of these scenes was encoded using an HRR as outlined in table 5 with Gaussian random vectors for each of the features. The model was trained on a sequence of 15 rooms and given a memory test after each room. We assumed that event boundaries corresponded to entering and exiting a room and constrained the model to infer an event boundary only at this time. The model was free to choose to reuse an event model or infer a new one for each room.

**Table 5.** Scene representation for Simulations of Radvansky and Copeland (2006). Each scene was composed of vectors corresponding to the features in the scene.

| Scene | Features |
| --- | --- |
| 1 | Enter + Current Room |
| 2 | Object A + Current Room |
| 3 | Object B + Current Room |
| 4 | Leave + Current Room |

The key manipulation in the simulations was the shift vs no-shift conditions, modeled as a change in event label memory precision for the ongoing event. As we alluded to above, event label memory precision was increased for the no-shift condition relative to the shift condition. The two conditions were otherwise identical. Error was assessed by the probability that a corrupted memory trace was included in the reconstruction sample. Mirroring the results in the human studies, the model had lower error in the no-shift condition than in the shift condition (Figure 13).

**Figure 13.**
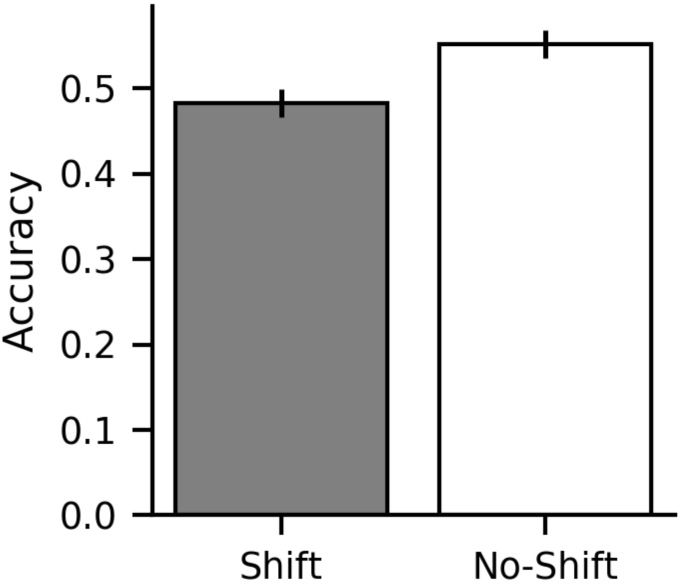
Simulations of the task in Radvansky and Copeland (2006). Reconstruction memory accuracy is shown for items both after an event boundary (*Shift*) and before an event boundary (*No-Shift*).

### Event boundaries improve overall recall

While these short-term memory effects suggest that event boundaries interfere with memory, the relationship between event structure and subsequent memory is not always intuitive. Overall, extracting relevant event structure tends to improve memory, as subjects with better segmentation judgments tend to have better subsequent recall (Sargent et al., 2013; Zacks et al., 2006). Similarly, studying a list of items or videotaped lectures in multiple contexts leads to better overall memory (Smith, 1982, 1984; Smith & Rothkopf, 1984). Within the context of SEM, event structure plays an important role in the reconstructive memory process. Poor segmentation leads to a noisier reconstruction process and thus worse overall memory.

Interestingly, the benefits of event boundaries within the reconstructive memory process extend to cases where the sequence of studied memory items are random and where there is no clear relationship between events and studied items. In a series of studies, Pettijohn and Radvansky (2016) demonstrated that when subjects were given a list of items separated by a physical or virtual event boundary, subsequent recall was higher overall. This suggests that segmentation itself influences memory and is important for overall memory irrespective of environmental statistics.

Here, we probe the model for these effects by simulating experiment 1 in Pettijohn et al. (2016). In this experiment, subjects were given a list of 40 words to remember while moving between four locations in physical space, divided into 4 ordered sub-lists of 10 words each. Subjects read one sub-list (10 words), then moved to a new location in space, either in a new room (*shift condition*) or a new space in the same room (*no-shift condition*) that was equated for physical distance and read a second sub-list (10 words). Subjects were then given a distractor task and finally asked to recall as many of the words as possible from both sub-lists. Subjects had higher recall accuracy in the shift condition than in the no-shift condition.

We simulated these effects by generating a list of 20 items, each as a Gaussian random vector. We trained the model on the list, either constraining the model to learn all items within a single event (no-shift) or assuming a single event boundary halfway through the list (shift). We then used the reconstruction procedure to create a reconstructed memory trace and probed memory recall as the probability that each corrupted memory item *ỹ_i_* is included in the reconstructed trace, defined by equation 14. Overall, the accuracy is higher in the switch than in the no-switch condition (Fig 14), replicating the findings of Pettijohn et al. (2016). During the reconstruction process, the uncertainty about a single item propagates to its neighbors as a consequence of the dynamics of the event schema. In general, the event schemata reduce the overall reconstruction uncertainty by regularizing the process, but they nonetheless propagate uncertainty between noisy scene memories. Because the dynamics end at boundaries, boundaries prevent the uncertainty from spreading.

**Figure 14.**
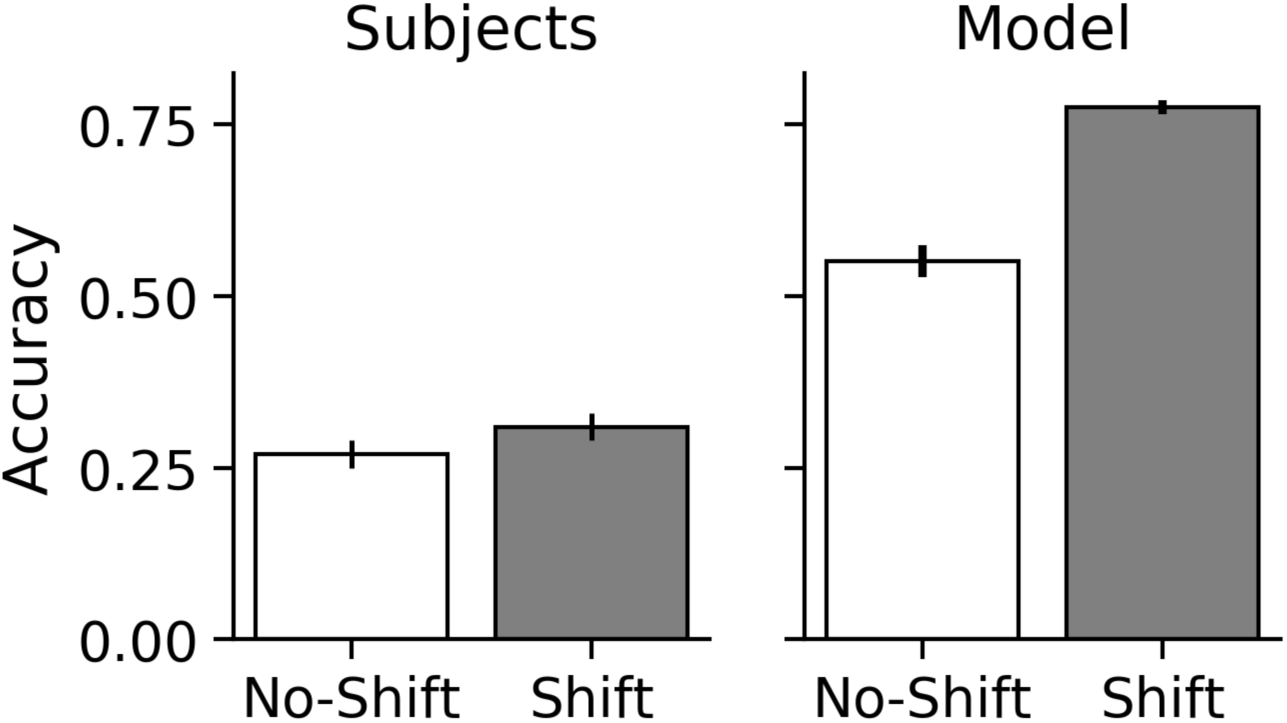
Simulations from Pettijohn et al. (2016). *Left*: Human subjects have higher recall in the shift vs no-shift conditions (data reproduced from Pettijohn et al., 2016). *Right*: Model has higher recall in shift vs no-shift condition.

### Sequential recall

Even as event boundaries improve memory performance overall, they introduce specific deficits to sequential recall. Given narrative texts that include a temporal shift, subjects are worse when remembering the next sentence immediately after a temporal shift than immediately before (Ezzyat & Davachi, 2011). This impairment of sequential order memory by boundaries occurs even in the absence of a naturalistic event structure (DuBrow & Davachi, 2013, 2016) and is not associated with an impairment of associative memory (Heusser et al., 2018). This suggests that the event structure that subjects learn (as opposed to the structure of the task) is responsible for this memory effect. In the context of the model, SEM continuously estimates the transition structure between scenes as the event dynamics, irrespective of the regularity or unreliability of that structure. Introducing a boundary disrupts learning an association between successive scenes. This becomes apparent in the reconstruction process, where the presence of boundaries disrupts sequential memory, even as they aid reconstruction overall.

To demonstrate this, we simulated experiment 1 of DuBrow and Davachi (2013). In the original experiment, subjects were presented with 400 items sequentially (200 celebrity faces and 200 nameable objects) across 16 study-test rounds of 25 images each, while performing a task in which subjects either made a male/female judgment (faces) or a bigger/smaller judgment (nameable objects). Following each round, subjects were asked to recall each item they saw in order. Sequential recall accuracy, as measured by direct transitions between consecutive items, was higher for items immediately before a task/category switch than immediately after.

We simulated 5 events, alternating between two categories after the presentation of 5 items each. Each item was embedded by combining an item-specific factor (i.e., a random Gaussian vector) and a shared category feature (itself a random Gaussian vector) with addition to encode the similarity relationship between items of the same category. As in the previous memory simulations, the model inferred a single event label for all of the scenes within an event and event boundary locations were provided to the model as a simplifying assumption of the task. As the deficit in serial recall can be interpreted as contradictory to the findings of Pettijohn and Radvansky (2016), we have re-used the same parameter values in both simulations to show that the same parameters can result in both behaviors. The reconstruction was performed as previously described, and SEM inferred a reconstructed memory trace and event label for each moment in time. Order memory was assessed by measuring the probability that two sequential probe items were reconstructed in the correct sequential order. Thus, if item *B* followed item *A* in the stimuli, it was scored as correct if it appeared in this order in the reconstruction, regardless of the position of these items in the reconstructed sequence.

Figure 15 summarizes the results of the model simulated on this task. As expected, the model has higher serial recall accuracy for items studied immediately prior to the category/task switch than after, mirroring previous empirical results. Mechanistically, this occurs because the model learns the dynamics over the scenes within an event, whereby each item is learned as a function of the previous item within the event, but there is no direct association between sequential items in separate events. This reflects the hypothesis that people are continually and automatically learning about temporal structure. The reconstruction process, which regularizes the corrupted memory trace with the learned dynamics, thus aids the acquisition of within-event sequence knowledge. Overall temporal information is not otherwise lost, as the identity of each event and the within-event dynamics help order the entire sequence, but there is less of an association between two sequential items across an event boundary in the reconstruction.

**Figure 15.**
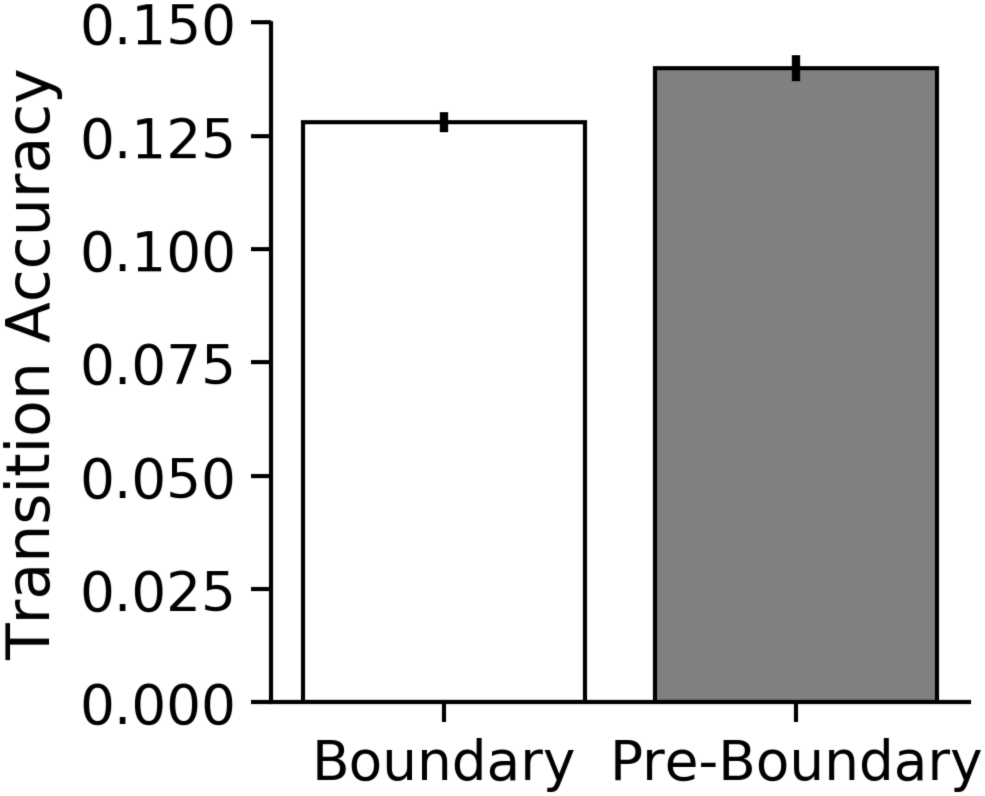
Serial recall across an event boundary. The proportion of correct transitions between an item and the following item was worse across an event boundary than immediately prior to the boundary.

One potential concern is whether these results are driven by our choice of parameter values. The temporal corruption noise parameter *b* is of particular interest because it directly controls how veridically order information is preserved in the noisy memory trace. Unsurprisingly, larger values of *b* lead to worse serial recall, but – more surprisingly – larger values of *b* also lead to better overall reconstruction (Appendix D, Figure 18). Intuitively, we might expect more temporal noise to increase the uncertainty of the reconstruction overall, but increasing temporal corruption loosens a constraint on the reconstruction process. This has the result that a corrupted memory trace is more likely to be included in the reconstructed memory, but at a cost of worse temporal order. As a comparison, the feature corruption noise parameter τ largely influences overall reconstruction and does not meaningfully influence order memory. Importantly, changes in the parameter value of *b* does not differentially affect serial recall in the boundary or pre-boundary trials (see Appendix D for details), indicating that the qualitative effect here is not driven by this parameter. This can be seen in Figure 16, which shows the order memory in the boundary and pre-boundary trials for 4 values of *b* but otherwise hold the parameters of the simulation the same.

**Figure 16.**
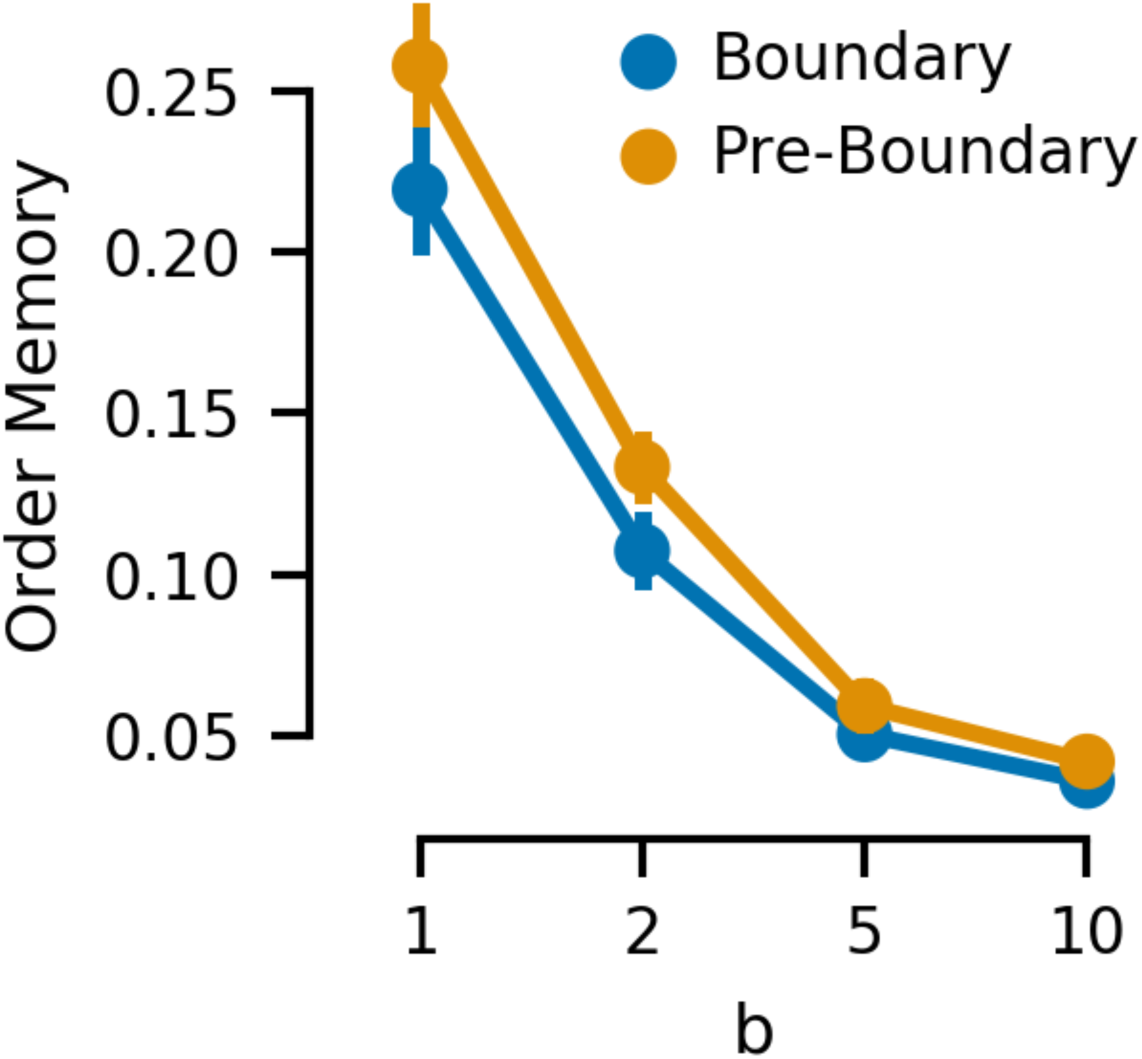
Sensitivity analysis for serial recall. Simulations of serial recall accuracy are shown for the boundary and pre-boundary trials for four values of temporal corruption noise *b*.

## Discussion

In this article, we have tackled the problem of learning and understanding events through the lens of probabilistic reasoning. We have made 3 main contributions with our model: first, we demonstrated that SEM can produce human-like segmentation of naturalistic data, suggesting that the computational principles outlined here are scalable to real-world problems; second, we have shown how events can capture generalizable structure that can be used for multiple cognitive functions; and third, we have shown that these principles are sufficient to explain a wide range of empirical phenomena.

### Segmentation

In the current work, we have built on previous models of event segmentation and shown that our model can produce human-like segmentation of naturalistic datasets. While the low-dimensional and often non-ecological stimuli used to evaluate previous computational models are diagnostic of computational principles, it is important to show these principles are sufficient to account for naturalistic environments. Without this demonstration, it is unclear whether the behavior produced by the model is a consequence of the artificiality of the stimuli. Furthermore, as we can show that human event boundaries mirror the boundaries predicted by the model, we can argue that human event segmentation is sensitive to the same underlying regularities in the stimuli as the model.

Previous computational models of events have, like SEM, used recurrent neural networks to learn the event dynamics. Even as the models differ in the specifics of the networks used, their similarity leads to qualitatively similar predictions. This is most evident in our simulations of community structure (Fig. 7). SEM recreates the same qualitative pattern of behavior as both the Schapiro et al. (2013) model and human subjects. All three show an increased tendency to delineate an event boundary at community transition points. It is not surprising that SEM and the Schapiro model learn similar event boundary points given the similar method of learning event dynamics. Nonetheless, these event boundaries in SEM correspond with increased prediction error, even as the Schapiro task was designed to *not* elicit prediction errors at community transition points. As we previously noted, this is a consequence of the distinction between the generative process of the task (in which community transitions are equally probable as other transitions) and what the agent learns about the task. SEM embodies a set of assumptions and inductive biases that differ from a direct inversion of the Schapiro task, and this leads to the experienced prediction errors at community boundaries.

Algorithmically, SEM is closely related to the model proposed by Reynolds et al. (2007). Both models employ a hierarchical process where event dynamics are learned with a recurrent neural network at the lower lever, and where a higher level *event* constrains the lower-level model. These models are also similar in that prediction errors play a key role in determining the identity of the higher-level *event* (Equation 6). Computationally, the type of gating mechanism used in the Reynolds model is similar to nonparametric clustering with a Chinese Restaurant Process prior (Collins & Frank, 2013), and while we make different commitments and assumptions, similar computational principles drive computation in both models. More broadly, using a recurrent neural network to learn temporal dependencies is a powerful computational tool, and has become more common recently in models of cognition. In a notable piece of recent theoretical work, J. X. Wang et al. (2018, 2016) argue that an architecture of stacked recurrent neural networks is sufficient to explain a host of empirical findings in human reinforcement learning and serves as a good model for the prefrontal cortex. Unlike SEM, their model requires extensive pre-training but nonetheless generalizes to novel task-variants efficiently. A similar architecture was also employed by Butz, Bilkey, Humaidan, Knott, and Otte (2019) to show that events are useful in goal-directed planning. They trained an agent to learn a forward model of states using recurrent neural networks, and showed that allowing these networks to be contextualized by events leads to more efficient learning in an artificial domain.

SEM is also related to computational techniques used to model sequential processes by inferring latent states. The hidden Markov model (HMM) is an instructive comparison because, like SEM, it assumes each observation is generated by a latent process associated with a discrete latent variable (Bishop, 2006; Rabiner, 1989). The key difference is that the HMM does not model the internal dynamics of each latent state; instead, HMMs assume that observations are generated by a stationary process conditioned on this latent state. While this limits the HMM as a cognitive model of events, it is nonetheless a useful empirical model in a variety of segmentation tasks. Goldwater, Griffiths and Johnson (2009; reviewed in Teh & Jordan, 2010) used an HMM with a uni-gram distribution over phonemes to segment words. Recent work by Baldassano et al. (2017) used HMMs to model event boundaries in human fMRI data, providing strong evidence that people spontaneously generate event boundaries with realistic experience. A more closely related family of models is the switching linear dynamical system (Fox, Sudderth, Jordan, & Willsky, 2010; Ghahramani & Hinton, 1996). These models are a generalization of the HMM and, like SEM, assume a dynamical process associated with each latent state. As the name implies, the model assumes a mixture of linear dynamical systems as its underlying process. This simplifying assumption facilitates learning the types of dynamical processes that constitute events, albeit at the cost of strong constraints on the form of these dynamical systems. This contrasts with SEM, which does not impose a linearity constraint, and can be viewed as a form of switching non-linear dynamical system. This distinction between linear vs non-linear dynamics is important for our choice of structured embedding space. SEM can learn an arbitrary sequence of bound scenes, a problem a linear model would struggle with.

### Generalization

A key difference between SEM and other computational models of event cognition is how SEM generalizes previously learned events to novel experiences. SEM accomplishes this via nonparametric Bayesian clustering, which allows SEM to learn and reuse events. Furthermore, because SEM assumes that events exist in a structured and distributed representational space, this generalization applies not only to the surface features of the event, but also to its underlying relational structure. As such, SEM can generalize an event dynamic even when many of the features are different, including role/filler bindings (Figure 9). The generalization of the relational structure is advantageous because its dynamics are thought to be smoother and thus easier to generalize than the surface features of a task (Radvansky & Zacks, 2011; Richmond & Zacks, 2017).

This places the burden of encoding structure on the representational space (in our case, an HRR), and allows the dynamics to be learned with a parameterized function. While we do not make a strong commitment to how this representational system is learned by neural systems, we note that the convolutions required to compute an HRR are thought to be plausible in biological networks (Eliasmith, 2013; Yamins & DiCarlo, 2016). Regardless of how this representational space is learned, smooth functions are sufficient to generalize because similar structures are represented with similar vectors. This allows SEM to leverage relational structure when identifying the event boundaries, and consequently learn more general event dynamics that abstract away some surface features. Hence, the event dynamics SEM learns are consistent with previous theoretical accounts in which events are encoded in terms of abstract, high-level features (Radvansky & Zacks, 2011).

The use of nonparametric clustering (i.e., the sticky-CRP) prior with the learned (and structured) dynamics produces novel empirical predictions. It might be unsurprising that people can learn an event schema and generalize the dynamics to a new set of role/filler bindings, but SEM makes predictions about when those dynamics should be generalized. Specifically, the model predicts that unpredictable dynamics or structure violations will drive scenes to be clustered into a new event, preventing generalization. For example, a sequence of scenes presented in a novel order will be assigned a new event. The model also predicts that the order in which events are initially experienced can influence how they are assigned to different event schema. Two instances of similar, but predictably dissociable, events can be assigned to the same event schema if they are learned close together in time. However, if one of the two events is well learned, this may not happen as the expected uncertainty of the event model (i.e., the learned value of *β* in Equation 5) is low, and any small deviation from the event model is sufficient to drive segmentation. To provide an intuitive example, if a person has a very regular morning commute, they might quickly notice changes in the traffic patterns caused by new construction and form a new event, whereas in other less practiced travel situations, they may ascribe variations in traffic conditions to more unpredictable processes.

Neural and behavioral measures tend to support the claim that events represent high-level information. A recent study by Baldassano, Hasson, and Norman (2018) found a neural representation for events that shared a high-level schematic structure but otherwise had very different features. In this passive task, subjects either watched a movie or listened to an audio narrative that followed a shared script, such as ordering food in a restaurant, but varied the actors, genre, and timing. Brain activity in the medial prefrontal cortex was highly predictive of the script and could be used to align the timing of events within the story, regardless of modality and other features. This suggests that subjects maintained a form of structural information about the films independent of their low-level features. More broadly, relational information is critical to memory (Cohen, Poldrack, & Eichenbaum, 1997), and thus we would expect to see it in event representations. This neural evidence is consistent with earlier memory studies in which subjects tended to lose lower-level descriptive details of narrative texts (e.g., the exact words used in order) while nonetheless maintaining an accurate high-level description of events (Kintsch, Welsch, Schmalhofer, & Zimny, 1990; Zwaan, 1994).

A limitation of our model is that it does not address the hierarchical nature of event representations. Events are thought to be composed sequentially from multiple smaller events in a temporal hierarchy (Radvansky & Zacks, 2011). To provide an intuitive example, if washing dishes reliably follows dinner, a model of event cognition should learn this relationship, potentially as a single, larger event, even as it may be adaptive to learn separate events for dinner and washing dishes. Behavioral evidence provides some support for hierarchical events, as events of different timescales have temporally aligned boundaries, consistent with what we would expect from a hierarchy (Zacks et al., 2001). Neural evidence complements this picture: Even in the absence of an explicit segmentation task, events of different timescales are discoverable in fMRI data (Baldassano et al., 2017) and memory representations of events vary in timescale along the long-axis of the hippocampus (Collin, Milivojevic, & Doeller, 2015). Additionally, there is some evidence that events of different timescales can be joined together through learning. When shown a conjunction of events in a film repeatedly, subjects are less likely to mark an event boundary between them, suggesting they may cohere smaller events into larger events (Avrahami & Kareev, 1994). Neural evidence also supports this hypothesis: in an fMRI study in which subjects passively viewed separate animated stories, activity patterns linked to distinct events in the medial prefrontal cortex and posterior hippocampus became more similar when these events were subsequently linked narratively (Milivojevic, Vicente-Grabovetsky, & Doeller, 2015). This is consistent with the proposal that people may learn smaller-scale events and combine them to form larger events in a hierarchy.

In the current model, we have not implemented a mechanism of learning hierarchical events. Nonetheless, there are two ways by which SEM could be extended to learning an event hierarchy. The first strategy is to learn events of a fixed timescale and directly learn the relationship between these events, either through transition dynamics between these events or through multiple levels of hierarchy. The sticky-CRP we use to model the transition dynamics does not encode this relationship, but it can be extended to do so in principle. The second strategy is to learn events of multiple different timescales in parallel. This is not a substantial modeling challenge for SEM, as longer or shorter events can be learned by scaling the prior over event noise parameter (*β* in equation 2), effectively changing each event’s sensitivity to prediction errors. To the degree that event boundaries are driven by environmental statistics, event boundaries from longer events naturally align with shorter events. These events of different timescales could be, in principle, combined in a mixture model for the purpose of making a unified forward prediction.

A further (and related) limitation of our model is that it is not compositional: elements of learned structure cannot be combined across events. This contrasts with human cognition, which is believed to be *systematic*, meaning that component pieces of knowledge can be combined in a rule-like manner to give rise to novel thoughts (Fodor & Pylyshyn, 1988; James, 1890). For example, an event representing a lunch meeting could, in theory, be composed of events separately representing lunches and meetings. This requires a different form of structure than the form outlined here, though it is worth noting that the false memory paradigm (Figure 12) does show aspects of a productive (or constructive) system, creating a novel trace composed from distinct pieces, without an explicit mechanism to do so.

To share structure between different events, these events need to be represented compositionally. In SEM as currently instantiated, events are learned as indivisible units with a single, learned function representing the dynamical structure. In principle, this dynamical structure can be decomposed and generalized independently between events. This may be adaptive because it simplifies the statistical problem posed by generalization (Franklin & Frank, 2018), and thus can simplify representational demands by a combinatorial factor, dramatically accelerating learning.

How event dynamics can be learned compositionally and generalized between events is an open question for future research. We can outline a few possible mechanisms for a compositional system. Perhaps the simplest form of compositionality would be to learn the dynamics of subsets of features independently and combine the independent predictions. As an intuitive example, the trajectory of a bird flying is generally independent of the trajectory of cars stopping at an intersection. If we observe both in the same scene, we might combine their previously learned patterns independently to generate a single prediction for the next scene. In the embedding space we’ve described, this is equivalent to linearly combining the predictions of two different systems via vector addition.

A second intriguing possibility coming from machine learning and function learning is to directly combine the predictions of multiple learned event models with composable kernel functions. Complex functions can often be decomposed into a combination of multiple, simpler functions (e.g., linear or periodic functions) with kernel methods (e.g., Gaussian processes or support vector machines; Duvenaud, Lloyd, Grosse, Tenenbaum, & Ghahramani, 2013; Schulz, Tenenbaum, Duvenaud, Speekenbrink, & Gershman, 2017). To the degree that we expect event dynamics to resemble combinations of multiple simpler dynamics (e.g., the zigzag line of a ship tacking in the wind resembles a combination of a linear and periodic function) then these compositions may provide a good model of how these components are learned.

An altogether different approach to decomposing the structure of individual events is to learn all events within a single, large event-model, for example in a neural network. Such a model, having never segregated the representation of different events, could potentially combine structure learned in different events naturally. Elman and McRae (2019) demonstrated this by training a neural network model on two different events, one in which a person cuts food in a restaurant with a knife and another where the same person cuts themselves with a knife and bleeds. They then gave the model an event in which the person was in the restaurant and cut themselves, and the model correctly inferred that the person bleeds, combining elements of structure from the two events. However, this approach comes at the cost of ignoring hierarchical event structure, which can lead to generalization impairments, as we show in Figure 11.

### Memory

The memory component of our model builds on several previous probabilistic models of memory. Most directly, it is an extension of the *Dirichlet process-Kalman filter* model (DP-KF) proposed by Gershman et al. (2014), a model of memory using a nonparametric form of the switching Kalman filter. In the DP-KF, observations were assumed to be generated by a switching linear-Gaussian diffusion process, and memory was modeled in terms of inference over this process. SEM is, in many ways, a similar model and can be seen as an extension of the DP-KF with learnable (as opposed to random diffusion) dynamics defining each event.

Most notably, the models are similar in how the partitioning of individual memory items into events (or clusters in the DP-KF) influences how they are smoothed in memory. In SEM, scenes are partitioned by event, and subsequent memory smoothing has the effect that scenes from the same event will influence each other more so than they will influence the smoothing of scenes in different events. Assuming that events are meaningful units, this is an efficient use of data corresponding to effectively pooling relevant sources of information and isolating irrelevant ones. Normatively, this can prevent memory interference by separating potentially competing stimuli, and is very similar to previous accounts of event boundaries in memory, in which boundaries prevent irrelevant information from spreading in a type of fan-effect (Radvansky, 2012; Radvansky & Zacks, 2017). Importantly, this interacts in the model with event dynamics, such that similar scenes experienced in multiple events will be separated in memory by their associated event models, and thus interference between the two will be lessened. Behaviorally, slowly drifting sensory stimuli are averaged together in memory tasks, whereas there is less averaging of stimuli across punctate change-points even in the absence of a context change, supporting a clustered organization of items in memory (Gershman et al., 2014).

Where SEM differs from the DP-KF and other accounts of memory is how the information from individual scenes is combined. In SEM, scenes associated with an event are smoothed with the regular trajectories of that event. This has the effect of regularizing memory traces towards an average event trajectory, not an average of all of the scenes, as a reduced model without dynamics would do. This mechanism differentiates SEM from prior models of memory. The degree to which memories are regularized toward an average event trajectory interacts with the inference of individual events. For example, events with small variations of a similar temporal structure will be more likely to be associated with the same event schema than events with dramatic differences. As a consequence, events with conserved structure are regularized to their schema-typical trajectory and events with a distinct temporal structure are not equivalently regularized. More concretely, familiar scenes that are presented in a surprising order will tend to be regularized *less* (and thus, closer to their original value but with higher overall noise) as they are likely to be assigned to a novel event with uncertain dynamics. Conversely, if an event is typical both in terms of content and temporal dynamics, it will be remembered better, but will also show more schema-consistent constructive inferences. This produces a systematic memory error in which schema-atypical scenes are regularized towards schema-consistent scenes. These mechanistic differences form the basis of empirically testable behavioral predictions: regularization can be measured by the degree to which reported memory differs from the original stimuli in the direction of the event dynamics.

In contrast to the latent structure modeled by both SEM and the DP-KF, the Temporal Context Model (TCM; Howard & Kahana, 2002) and the related Context Maintenance and Retrieval model (Polyn et al., 2009) have proposed that memory is encoded in an evolving temporal context. In TCM, a context vector that encodes an average of recently seen items is bound to each new memory item. This context drifts over time as new percepts are used to update it, and is bound in memory to individual items. The Context Maintenance and Retrieval model adds to this the property that task changes induce large context shifts, which partitions traces from different contexts. These models capture the phenomenon that when asked to recall lists of items, recall can be temporally organized even when participants are not constrained to recall items in order (Kahana, 1996), a property of behavior captured by TCM (Howard & Kahana, 2002). In contrast to a constantly drifting process, SEM models sequential dependencies as dynamical systems, and uses a recurrent neural network to learn them.

Extant empirical evidence suggests that neither abrupt shifts nor slowly drifting memory contexts are sufficient accounts of memory (DuBrow, Rouhani, Niv, & Norman, 2017). SEM offers a potential solution by embodying both. Nonetheless, it is not always clear whether an individual experiment requires either clustered structure or event dynamics as they sometimes produce similar results. To dissociate this experimentally, these two forms of statistical structure need to be both present and decorrelated in a task, ideally through direct manipulation. We have shown how these two forms of structure produce different patterns of generalization but demonstrating how the two forms of structure interact to influence memory within a single task is a question for future empirical research.

Our memory model shares several features with other memory models. SEM is conceptually similar to the *schema-copy plus tag* model, in which a memory trace is composed of a corrupted copy of a schema and a set of schema atypical features (Graesser & Nakamura, 1982). The perspective of memory retrieval as inference over a noisy memory trace is shared with the REM model (Shiffrin & Steyvers, 1997), and the regularizing effect of the event dynamics is qualitatively similar to the category bias proposed by Huttenlocher et al. (1991).

SEM offers a novel and complementary approach to reconstructive memory that is meaningfully different from prior memory models. Broadly, reconstructive memory can be thought of as the process by which we reconstruct the past from fragmentary recollections. Neisser (1967) made a famous analogy about how reconstructive memory is like a paleontologist assembling a dinosaur from dug-up bones. Many existing models (e.g., Norman & O’Reilly, 2003; Raaijmakers & Shiffrin, 1980; Shiffrin & Steyvers, 1997) are concerned with which bones are found (how a memory fragment is retrieved), whereas with SEM we take the perspective of the paleontologist and ask how do we put the pieces together.

The difference between the reconstruction process of SEM and the generation of memory traces of prior models is similar, but not equivalent, to the distinction between semantic knowledge and episodic memory. The noisy memory trace SEM uses in the reconstruction process can roughly be thought of as a form of episodic memory, whereas the event dynamics are more akin to semantic knowledge used to organize them during reconstruction. This comparison breaks down when we consider that novel events in the model have episodic-like qualities; event dynamics for a newly learned event capture much more detail about the specific event than would be expected for an event that has been experienced multiple times. As an event is experienced multiple times, the learned dynamics in SEM come to reflect the more generalized dynamics, averaging away the specific features of an individual event. This leads to a transition of an episodic-like event dynamic to a semantic-like event over time.

Likewise, SEM does not explicitly differentiate between types (a general class of symbol) and tokens (a specific instance of that symbol) but does encode knowledge of both in memory. A bound scene with fillers has both type and token information of individual symbols. In SEM, an event model that has experienced the same structure multiple times with different fillers will average over the fillers, which can be interpreted as type (i.e., semantic) knowledge of those symbols. Noisy memory traces used in the reconstruction process have information about both, albeit in corrupted form. For well-learned events, the reconstruction process will tend to amplify the type information, as that is what the event model has learned, but the final trace will preserve both type and token information.

### Neural Correlates

A further open question is how our model of events is implemented in the brain. A growing body of evidence has linked the hippocampus and the posterior medial network to event segmentation and memory (Ranganath & Ritchey, 2012). The hippocampus plays a crucial role in binding features and contextual information (Davachi, 2006; Diana, Yonelinas, & Ranganath, 2007; Eacott & Gaffan, 2005; Eichenbaum, Yonelinas, & Ranganath, 2007; Knierim, Lee, & Hargreaves, 2006; Ranganath, 2010), and could therefore play a role in binding features of an event. Consistent with this hypothesis, recognizing objects across an event boundary is associated with an increase of hippocampal activity (Swallow et al., 2011), and event boundaries themselves are linked to an increase in hippocampal signal (Baldassano et al., 2017). The anterior hippocampus also appears to play a role in representing events; its representation is sensitive to temporal community structure present in a segmentation task (Schapiro et al., 2016).

The posterior medial network (PMN), a component of the default network, has anatomical connections to the hippocampus and encompasses the parahippocampal, posterior cingulate, retrosplenial, ventromedial prefrontal cortex, precuneus and the angular gyrus (Ranganath & Ritchey, 2012). Hippocampal and PMN regions consistently co-activate during recall of experienced events, recollection of context, retrieval of temporal sequences, imagination of future events, and spatial navigation (Ranganath & Ritchey, 2012; Rugg & Vilberg, 2013; Spreng & Grady, 2010). A linking thread through these tasks is the need for a constructed event representation.

Consistent with this idea, fMRI and electrocorticography studies have shown that regions of the PMN network integrate over long timescales (hundreds of seconds; Hasson, Chen, & Honey, 2015), and PMN activity is modulated by event boundaries (Kurby & Zacks, 2008). Furthermore, event boundaries are identifiable in the angular gyrus and posterior medial cortex in unsupervised fMRI analysis (Baldassano et al., 2017). In terms of memory, medial prefrontal cortex brain activity has been associated with schema-consistent memory (van Kesteren et al., 2013), further implicating the posterior medial network in event processing.

Integrating across these findings, we propose that event dynamics could be learned and represented by the PMN (Figure 17). We further propose that the ventromedial prefrontal cortex (vmPFC) encodes the posterior probability distribution over event models, as the vmPFC has been previously implicated in latent cause inference (Chan, Niv, & Norman, 2016) and state representation (Schuck, Cai, Wilson, & Niv, 2016; Wilson, Takahashi, Schoenbaum, & Niv, 2014), two similar computational problems. Finally, we hypothesize that the PM-hippocampal interactions are responsible for the reconstructive memory process, as this process involved the re-instantiation of scene dynamics with memory traces.

**Figure 17.**
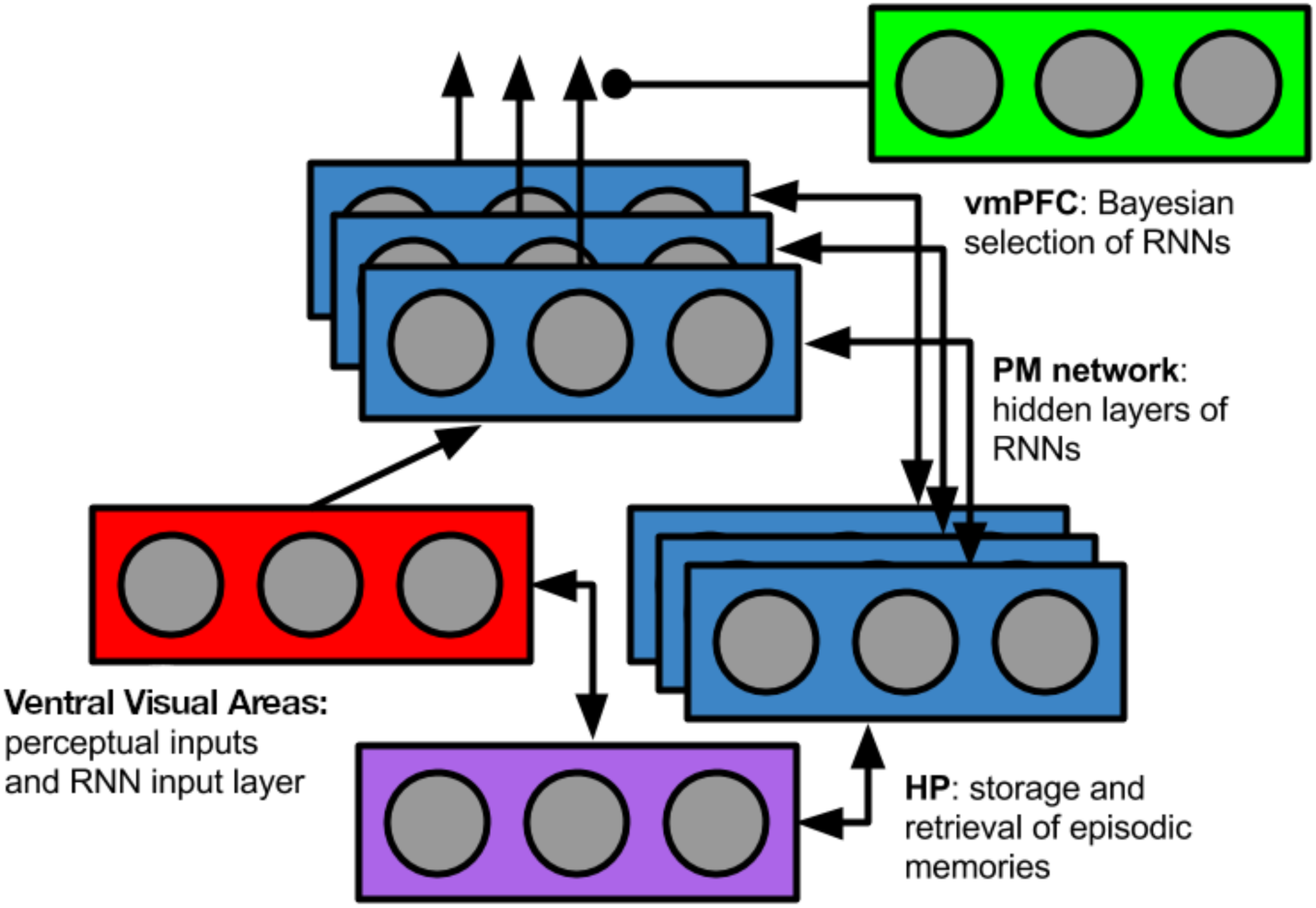
Neural Correlates of SEM. Diagram of SEM architecture with corresponding brain regions. Information about entities from the ventral visual areas is fed into event models, which are instantiated in the posterior medial network (PMN) (blue) as recurrent neural networks (RNNs). vmPFC (green) is hypothesized to select the currently relevant event schema/RNN. The hippocampus is hypothesized to support storage and retrieval of event-specific information (i.e., activity patterns in the currently-selected RNN).

**Figure 18.**
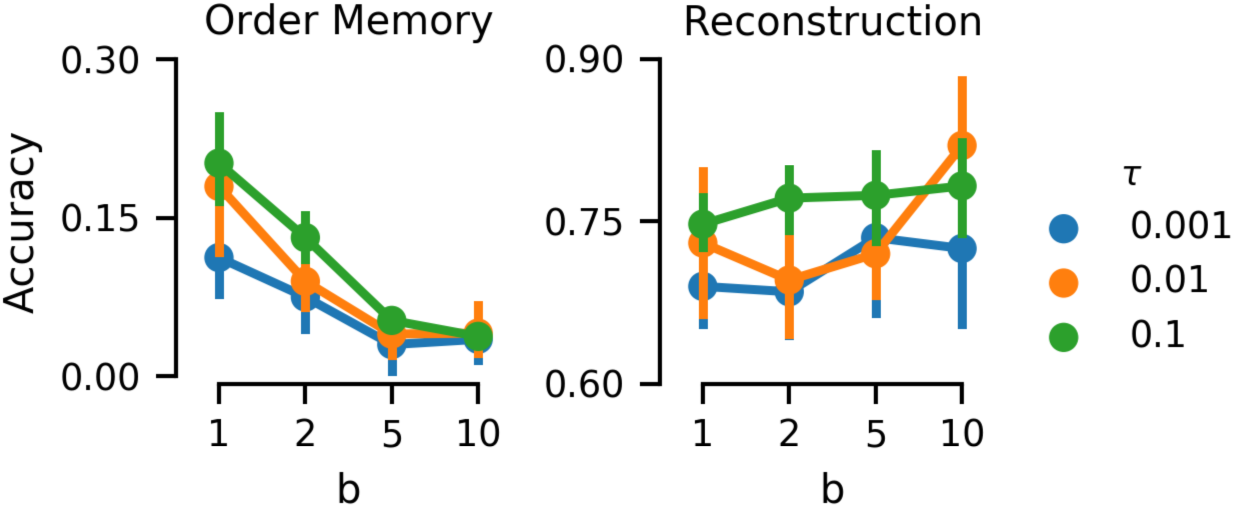
Sensitivity analysis of memory parameters. Order memory (*left*) and overall reconstruction accuracy (*right*) are shown for different values of the temporal corruption noise (*b*) and feature corruption (τ). Higher values of *b* decrease order memory but aid overall reconstruction. Increased values of τ correspond to both improved order memory and increased reconstruction accuracy.

Hence, we would expect to see these regions modulated by events in different ways as a function of the experimental paradigm. For example, we would expect to see stronger modulation of the vmPFC in tasks where the strength of evidence for each individual event model varies dramatically. Likewise, we might expect to see modulation in the hippocampus and PMN in tasks that manipulate event reconstruction but less modulation of vmPFC in such a task. We also might expect the relationship between event segmentation and subsequent memory measures to increase following sleep, reflecting prior research linking sleep and the increased semanticization of memory (Dudai, Karni, & Born, 2015), and we would expect this to be reflected by hippocampal replay. Although all of these predictions are strongly motivated by the existing literature, our neural predictions are an open empirical question for future research.

A related issue is how the computational-level model relates to a circuit-level implementation. We note that many key aspects of our theoretical account are neurally plausible. Of primary interest is how event dynamics are learned by the PMN. One component of this problem is learning an effective representational space, which can dramatically simplify the problem of learning event dynamics. While the exact representational format of scenes is an open question, Richmond and Zacks (2017) argued that we should expect these representations to be smooth over time, as event dynamics are consequently easier to learn and generalize. This mirrors techniques in statistics and machine learning, where the transformation from a complex representational space to a simpler, intermediate representation is a commonly used tool (Bishop, 2006).

To learn the dynamics over these representations, we have used recurrent neural networks as a function approximator. Gating mechanisms, the ability to selectively store the internal state of the network, are a critical element of these networks and make them well-suited to learning sequential dependencies (Hochreiter & Schmidhuber, 1997; LeCun et al., 2015). Gating mechanisms are common in biologically inspired models of human rule learning and reinforcement learning (Collins & Frank, 2013; Kriete, Noelle, Cohen, & O’Reilly, 2013; O’Reilly & Frank, 2006; Rougier, Noelle, Braver, Cohen, & O’Reilly, 2005; J. X. Wang et al., 2018), and are hypothesized to be supported by the midbrain dopamine system (Collins & Frank, 2013; Frank & Badre, 2011; O’Reilly & Frank, 2006). In line with these models, one possibility is that the midbrain dopamine system (including the ventral tegmental area and substantia nigra) provides an error signal for learning event dynamics through a gating mechanism similar to the ones found in artificial neural networks. This fits with the generalized view of dopamine prediction errors recently espoused by Gardner, Schoenbaum, and Gershman (2018). As an alternative possibility, the PMN may learn the event dynamics directly through local connectivity and its own prediction errors. How this may occur is an open question, but prior computational modeling work has suggested that gating mechanisms are consistent with cortical microcircuits (Costa, Assael, Shillingford, de Freitas, & Vogels, 2017), and that neural oscillations may be sufficient to generate a training signal (O’Reilly, Wyatte, & Rohrlich, 2014).

### Language and other outstanding questions

In the current work, we have not considered tasks that rely on the comprehension of natural language texts. In the event cognition literature, many of the effects that have been observed use narrative texts in their experimental design and measure reading speeds (for review, see Radvansky & Zacks, 2014). For example, subjects show slower reading speeds following a change in goal (Suh & Trabasso, 1993) or cause (Zwaan, Langston, & Graesser, 1995) in narrative texts, an effect that has previously been interpreted as reflecting an event boundary (Radvansky & Zacks, 2014). Memory for specific sentences read in a narrative declines across time even as the memory of the events described remains stable (Fletcher & Chrysler, 1990; Kintsch et al., 1990; Schmalhofer & Glavanov, 1986). In principle, SEM could be applied to natural language texts as long as these texts could be encoded into the logical scene description language that we embed into vector space using HRRs.

More broadly, while our model of events included the ability to encode structured representations, the degree to which it encodes semantic information is a function of the embedding space. This space is assumed as an input to the model, and when it is semantically meaningful, the model can generalize this semantic meaning to novel events through learned dynamics (Figure 9). However, when we only provide the model with an unstructured embedding space, it cannot produce variable bindings through event segmentation or with event dynamics alone. For example, in our simulations on the video data, an unstructured embedding of each of the videos was created using a variational auto-encoder. Our purpose was to demonstrate that SEM could generate human-like segmentation from pixel-level data via unsupervised training alone and without access to hand-tuned representations of objects. Nonetheless, these scenes contained objects in them, and a lack of a structured representation of these scenes likely made the segmentation problem more challenging. This is likely a larger problem in more complex environments. Each of the videos that we examined was simple, concerning a single person completing an everyday action. We suspect that, for example, a video in which several actors cooperate on a task, or a video with background motion (cars in the distance, birds moving around) would be much more difficult for the computational model without a structured representation. The question of how to develop structured representations of scenes is an ongoing question of research in computer science (see Herath, Harandi, & Porikli, 2017, for review), and progress in this area will aid our ability to model naturalistic datasets going forward.

## Appendix A: Variational Auto-encoder

A variational auto-encoder (Doersch, 2016; Kingma & Welling, 2013) is a dimensionality reduction technique that employs two networks, an *encoder* network to project the data into a lower-dimensional embedding space and a *decoder* network to re-project it into the original space. A Gaussian prior is defined over the embedding space and the network is trained to minimize the difference between the original image and its reconstruction, subject to the embedding prior. We used a variant of the Deep Convolutional Generative Adversarial Networks (DCGAN) architecture (Radford et al., 2015) for our encoder and decoder networks and use a maximum mean discrepancy kernel as a prior over our embedding space (Zhao, Song, & Ermon, 2017). Our encoder network consisted of a five layers: three convolutional layers (with 64×3, 128×3 and 256×3 channels, respectively) and two fully connected layers. All of the layers used a leaky, rectified linear activation function except the last layer, which was linear and 100-dimensional. Our decoder network reversed this process with a symmetrical network. Prior to training, each frame was downsampled to a resolution of 64×64 using linear interpolation. Randomly selected batches of frames were used with the ADAM optimization algorithm (Kingma & Ba, 2014) to train the autoencoder. A PyTorch implementation of our neural network is available at https://github.com/ProjectSEM/VAE-video.

## Appendix B: Holographic reduced representation

Holographic reduced representations (HRR) leaves intact the similarity structure of the composed vectors. HRRs consist of two operations, vector addition and circular convolution. Vector addition preserves similarity, such that if **a**, **b** and **c** are vectors, then then (**a** + **c**) and (**b** + **c**) are typically more similar to each other than **a** and **b** are to each other. This is always true for zero-mean orthogonal vectors and is true in the expectation for zero-mean random vectors in high dimensional space (e.g., **x** ∼ N (0*, I*)) To demonstrate this, compare the orthogonal vectors **a**, **b** and **c** with dot product as our measure of vector similarity, then **a**^T^**b** = 0 while (**a** + **c**)^T^(**b** + **c**) = **c**^T^**c**. Likewise, if we assume the two vectors **x** and **y** are similar to each other such that **x**^T^**y** ∕= 0, then adding orthogonal features to them does not change their similarity. If **x** and **y** are also orthogonal to **a** and **b**, then (**x** + **a**)^T^(**y** + **b**) = **x**^T^**y**. Thus, adding orthogonal feature vectors does not change the similarity between two vectors.

Using circular convolution as a binding operation also preserves the similarity structure. That is, if two vectors are similar to each other, then their convolution with a third vector will partially retain that similarity. We can show this by approximating a circular convolution with a tensor product (Doumas & Hummel, 2005; Plate, 1995), and noting that tensor operations are distributive, such that (**x** + **y**) ⊗ **z** = **x** ⊗ **z** + **y** ⊗ **z**. If two vectors **a** and **b** share a common factor **c** such that **a** = **a**_0_ + **c**, and **b** = **b**_0_ + **c**, where **a**_0_, **b**_0_ and **c** are orthogonal vectors, we can decompose their tensor product with **d** into the sum of two separate tensors

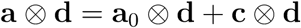

and

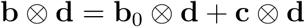

provided that **d** is orthogonal to **a** and **b**. Because both tensor products share a common tensor as a linear factor, we can use the arguments above to show that they are similar to each other. Thus, taking the tensor product of two similar vectors and a third random vector will result in a similar tensor product. Since circular convolution is a compressed tensor product operation (Doumas & Hummel, 2005; Plate, 1995), this argument will hold approximately for HRRs as well.

## Appendix C: Sampling reconstruction memory model

In order to sample the reconstructive memory model, we define a Gibbs sampling algorithm. The algorithm has three interlocking pieces: (1) sampling reconstructed estimates of the scene features 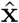 conditioned on samples of *w* and *ê*, (2) sampling estimates of *ŷ* from conditioned on samples of 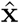 and *ê* and (3) sampling estimates of *ê* conditioned on samples of 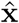 and *ŷ*. As each component is conditionally independent of the other, they can be iteratively sampled until convergence.

To initialize, at each time *n* we either draw a sample of 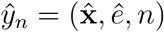 from either *ỹ* without replacement or assign it to be *y*_0_. The features of the scenes are initialized from a normal distribution and the sequence of events *e* is initialized from the sCRP prior.

### **Sampling** Pr(**x**|*ỹ, e*)

In the first step of the sampling algorithm, we draw samples of the scene features conditioned on the corrupted memory trace *ỹ* and the event label *e*. The probability of the feature vector **x**’ occurring at time *n* can be recursively defined under the generative model as:

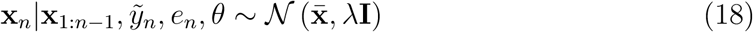

where the mean 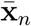 is a precision-weighted linear combination of the scene features of the memory trace 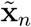 from the corrupted memory trace *ỹ_n_* and the predicted location of the embedded scene under the event model:

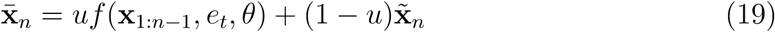

where *u* = *β*^−1^*/* (*β*^−1^ + τ^−1^) is the weight placed on the event schema prior, and λ = 1*/*(*β*^−1^ + τ^−1^) is the posterior variance.

### **Sampling** Pr(*ỹ*|**x***, e*)

The samples of the features vectors are then used to sample *ỹ* and *e*. The corrupted memory traces *ỹ* are sampled to determine both the time they occur as well as whether they are recalled at all. Samples of *ỹ* are drawn from the conditional distribution of the memory trace given the reconstructed features and event labels, which is defined by inverting the corruption processes (Equations 11, 12 and 13):

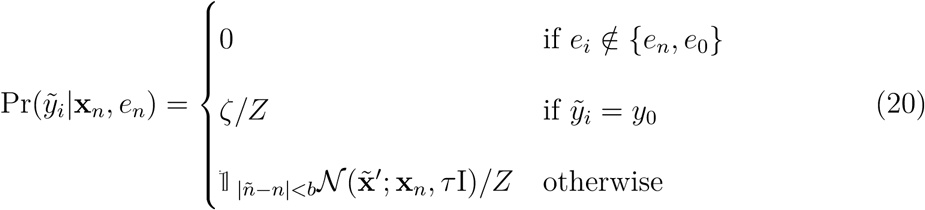

where 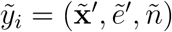, and where 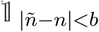 is an indicator function that returns 1 if |*ñ* − *n*| *< b* and 0 otherwise. Thus, the probability the corrupted memory item occurs at time *n* is zero if the corrupted event label is mismatched, or if the corr upted time index *ñ* is more than *b* steps away from *n*.

It is important to note that the probability of sampling a null event under this process is both a function of ζ and the estimate of **x***_n_*. This is because the normalizing constant

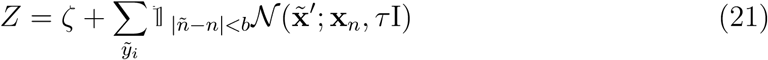

will typically be larger as a function of 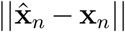, thus lowering the probability of a *null* event. Consequently, the quality of the reconstructed features will influence the degree to which a *null* token is included. It is also worth noting that the use of a uniform prior over a restricted range greatly simplifies the sampling problem when compared to, for example, a discretized normal distribution. Because we assume each memory token *ỹ_i_* is sampled without replacement, there are less than 2*N* ^2^*^b^*^+1^ valid assignments of *ŷ*, whereas an unbounded time corruption process leads to 2*N* ! valid permutations.

### **Sampling** Pr(*e*|**x***, ỹ*)

To complete the inference process, the samples of 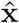 and *ŷ* are used to update the estimate of the sequence of events. Conditioned on **x**, *f* and θ, the probability of events *e*_1:_*_n_* under the generative process is recursively defined as:

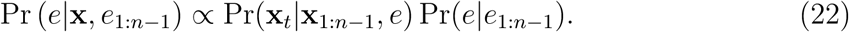

As previously noted, we store a corrupted memory of the event label *ẽ_i_* in the memory trace *ỹ_i_*. Conditioning on this memory trace, the reconstruction process is thus:

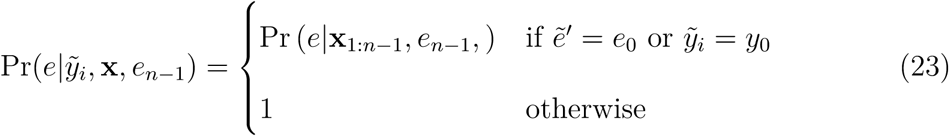

If the associated event memory *ẽ*’ is equal to the *null* token *e*_0_, or if *ỹ_i_* is the *null* memory item *y*_0_, then the generative process (Equation 22) is sampled. Otherwise, the memory label is taken from the estimated memory trace *ŷ_n_* In the full Gibbs sampler, we draw alternatively draw samples of 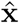, *ŷ* and *ê* until convergence. To this sample, we can apply different scoring rules to simulate different memory measures. For example, we model recall by whether a trace *ỹ* is in the final sample. These measures are specific to each simulation and we present each with the relevant simulation.

## Appendix D: Sensitivity Analysis of DuBrow and Davachi (2013, 2016)

To evaluate the sensitivity of the model simulations for the task presented in DuBrow and Davachi (2013, 2016), we simulated a single, constant run of the task with multiple values of memory corruption noise *t* and *b*. Using the parameter values *t* = [0.001, 0.01, 0.1] and *b* = [1, 2, 3, 4], 8 batches were simulated for each parameter. All of the other parameters were held constant as reported in Table 1. We assessed the effect of these parameters on overall order memory (the probability each item was reconstructed following the item that preceded it in the training data) and the overall reconstruction accuracy (the probability that the memory item was in the reconstruction sample). This was done using Bayesian logistic regression:

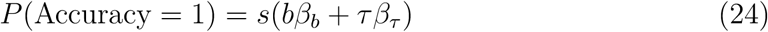

where *s*(*x*) = 1*/*(1 + *e*^−*x*^) and *β* is a regression weight. We also assessed whether there is a difference in order accuracy in the pre-boundary and post-boundary trials as a function of *b* by fitting separate regression weights for the two conditions with the following model and comparing the regression weights. We report the 95% highest density posterior interval (HDI; Kruschke, 2014)) and consider statistically significant effects as those not containing zero in this interval.

Higher values of *b* were found to increase overall reconstruction accuracy (*β_b_*: HDI=[0.14, 0.23]) and decrease order memory (*β_b_*: HDI=[−1.53, −1.15]). Increased value of τ did not lead to a change in order memory (*beta*_τ_: 95% HDI=[-2.41, 4.76]) but lead to higher reconstruction accuracy (*β*_τ_: 95% HDI=[5.49,12.11]). Critically, the effect of the value of *b* did not vary for the pre-boundary and post-boundary conditions (*β_b_pre__* − *β_b_post__* 95% HDI=[-0.58, 0.63]), suggesting the parameter values did not drive the qualitative pattern we report.

We further verified this result by simulating 12 batches of the full (DuBrow & Davachi, 2013, 2016) task for the values of *b* = [1, 2, 5, 10] and with τ = 0.01. For all of these values of *b*, pre-boundary serial recall was higher than boundary serial recall (Figure 16).

## Footnotes

1 For an explanation of this culinary metaphor, see Gershman and Blei (2012)

2 We could define prediction error using another similarity metric (e.g., angle cosine). With an alternate definition, we would nonetheless expect prediction error to negatively correlate with the log-likelihood of scene transitions, but this is not an axiomatic property of the model.

3 Implicitly, this corruption process assumes a uniform prior over features, Pr(**x**) ∝ 1. Alternatively, we could assume a generative process **x** ∼ *N* (0, ζI), but this will lead to a similar result so long as the norm of corrupted memory traces are similar, and ζ would likewise parameterize the degree of forgetting in the model for reasons discussed below.

4 A Hamiltonian path is a path through the graph that visits each node exactly once.

5 Formally, this is defined with the density function 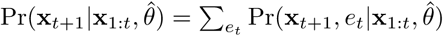.

6 We can interpret this structure linguistically, but we do not have to – this symbolic representation is valid whether we are watching the event occur or reading about it in a text.

7 This was done by setting the stickiness parameter λ and the concentration parameter *α* of the sticky-CRP to 10^6^ and 10^−6^, respectively, effectively forcing the model to treat scenes as belonging to the same event. Importantly, this manipulation does not affect the training of the event dynamics.

